# Precision Autism: Genomic Stratification of Disorders Making Up the Broad Spectrum May Demystify its “Epidemic Rates”

**DOI:** 10.1101/2021.07.28.454081

**Authors:** Elizabeth B Torres

**Affiliations:** Rutgers the State University of New Jersey, Psychology Dept; Rutgers University Center for Biomedicine Imaging and Modeling, Computer Science Dept.; Rutgers University Center for Cognitive Science

**Keywords:** Autism, genes, tissues, stratification, neurodevelopment, neurological disorders, neuropsychiatric disorders

## Abstract

In the last decade, Autism has broadened and often shifted its diagnostics criter a, allowing several neuropsychiatric and neurological disorders of known etiology. This has resulted in a highly heterogeneous spectrum with apparent exponential rates in prevalence. I ask if it is possible to leverage existing genetic information about those disorders making up Autism today and use it to stratify this spectrum. To that end, I combine genes linked to Autism in the SFARI database and genomic information from the DisGeNet portal on 25 diseases, inclusive of non-neurological ones. I use the GTEx data on genes’ expression on 54 human tissues and ask if there are overlapping genes across those associated to these diseases and those from SFARI-Autism. I find a compact set of genes across all brain-disorders which express highly in tissues fundamental for somatic-sensory-motor function, self-regulation, memory, and cognition. Then, I offer a new stratification that provides a distance-based orderly clustering into possible Autism subtypes, amenable to design personalized targeted therapies within the framework of Precision Medicine. I conclude that viewing Autism through this physiological (Precision) lens, rather than viewing it exclusively from a psychological behavioral construct, may make it a more manageable condition and dispel the Autism epidemic myth.

## 1. Introduction

According to the CDC, in the span of 16 years, the US moved from 6.7/1000 to 18.5/1000 autistics in the population of school age children [1]. Reportedly, this increase continues to move along an exponential rate, while maintaining a near 5:1 males-to-females ratio [2, 3]. This ratio prevents researchers from spontaneously reaching statistical power in any random draw of the population, when attempting to characterize the autistic female phenotype. Yet, motor features derived from endogenous neural signals in motor patterns, do identify the female phenotype [4–7]. This is the case even when digitizing the current clinical criteria that would otherwise miss females because of exclusive reliance on external observation [8, 9]. Likewise, subtle cultural biases built into the social-appropriateness criteria of the current instruments skew identification of underserved populations [1]. Consequentially, current interventions are far from being inclusive, or advocating for neurodiversity in the clinical data driving best-practices and evidence-based criteria for treatment recommendation [10, 11].

Despite sparse sampling in certain sectors of society, the shifts in diagnostic criteria have significantly broadened the detection rates to include now children with sensory issues and to allow comorbidity with ADHD under the Diagnostic Statistical Manual (DSM-5) [11]. This inclusion of ADHD in ASD contrasts with the former DSM-IV criteria, which would not allow comorbidities of ASD and ADHD, nor would it recognize sensory issues in Autism.

The challenges that broadening the diagnostics criteria bring to the science and practices of Autism are manifold [12], albeit discouragement by some clinicians from trying to stratify the spectrum into subtypes [13, 14]. Under current standards, motor, kinesthetic sensing, and vestibular issues are not part of the core symptoms of the original diagnosis in the DSM. These criteria also remain absent from psychological instruments like the ADOS test, currently used to diagnose different age groups [10, 15, 16]. However, kinesthetic sensing and motor/vestibular issues define several of the many disorders that today received the Autism diagnosis [17]. Among them are Cerebral Palsy [18, 19], Dystonia [20, 21], Tourette’s Syndrome [22] and obsessive-compulsive disorders (OCD) thought to be related to ADHD [23]. Besides these neurological disorders in ASD [24], others of known genetic origins enter in the broad criteria for Autism. Among them, various types of Ataxias [25] and Fragile X [26, 27] make up for a large percentage of individuals with Autism today. Despite profound physiological, systemic alterations and somatic-sensory-motor differences, these individuals will very likely go on to receive blanket-style behavioral modification-treatments-for-all, during early interventions. Furthermore, these behavioral modification interventions in the US, will continue later at the school, through the individualized education plan. Such treatments neither recognize, nor address individual phenotypic physiological features of these disorders of known genetic origins that, nevertheless, do enter in the Autism spectrum today.

Phenotypically, these disorders that currently also go on to receive the Autism diagnosis, are precisely defined by somatic, sensory-motor issues [28, 29] that manifest throughout the lifespan [30]. Their definition in their fields of origin, is nevertheless at odds with the current clinical “gold standard” criteria. In the DSM-5 [11] we read “*Hyper- or hyporeactivity to sensory input or unusual interests in sensory aspects of the environment (e.g., apparent indifference to pain/temperature, adverse response to specific sounds or textures, excessive smelling or touching of objects, visual fascination with lights or movement)*.” And in the DSM-5 criteria, motor issues are excluded owing to the confounds of symptoms induced by psychotropic medication, “*Medication-induced movement disorders are included in Section II because of their frequent importance in 1) the management by medication of mental disorders or other medical conditions and 2) the differential diagnosis of mental disorders (e.g., anxiety disorder versus neuroleptic-induced akathisia; malignant catatonia versus neuroleptic malignant syndrome). Although these movement disorders are labeled ‘medication induced’, it is often difficult to establish the causal relationship between medication exposure and the development of the movement disorder, especially because some of these movement disorders also occur in the absence of medication exposure. The conditions and problems listed in this chapter are not mental disorders.*” This neglecting of motor issues is enforced despite scientific evidence that even without medication, there are profound motor issues in Autism [5] that intensify with aging [30].

Further sidelining sensory-motor issues in Autism, within the ADOS booklet [10], under the guidelines for selecting a module, we read “*Note that the ADOS-2 was developed for and standardized using populations of children and adults without significant sensory and motor impairments. Standardized use of any ADOS-2 module presumes that the individual can walk independently and is free of visual or hearing impairments that could potentially interfere with use of the materials or participation in specific tasks*”.

Despite these caveats explicitly stated on their manuals, children with profound and highly visible motor, kinesthetic sensing and vestibular issues [31] go on to receive the Autism diagnosis that places them on a behavioral modification therapeutic pipeline that does not consider the brain-body physiology [32]. Clearly, there is a contradiction between the somatic-sensory-motor medical-physiological criteria and the social-appropriateness behavioral-psychological criteria explicitly denying the former. *Which one is it? And why are these important medical-physiological factors deemed secondary or co-morbid, when they are at the core of the basic building blocks necessary to develop and mainsocial behaviors?*

To better understand this tension between psychological criteria (dominating Autism research, diagnostics, therapies and services) and medical-physiological issues reported by peer-reviewed science [33], I here examine the genes linked to these neurological and neuropsychiatric disorders making up a large portion of the autistic population today and manifesting profound somatic and sensory-motor differences.

I investigate the pool of genes linked to a purely behavioral diagnosis of Autism attained through instruments that precisely sideline the somatic sensory-motor physiology (*i.e*., the ADOS/DSM-5). Specifically, the ADOS-2 research criteria inform the studies that support the confidence scores of the Autism-linked genes hosted by the Simons Foundation Research Initiative (SFARI). I leverage this research-based data repository of Autism-linked genes and use it as reference to compare its gene pool to the genes from other sources identifying neurological and neuropsychiatric disorders making up the Autism spectrum today. Given that those other disorders of known genetic origin are visibly affected by somatic-sensory-motor differences, but also receive the Autism diagnosis, I here ask if the gene pool of those disorders could help us stratify the broad spectrum of Autism into subtypes.

Stratifying the broad spectrum of autism based on available genetic information, would help us advance at least two areas of intervention. At the non-drug intervention level, if we were to learn that a subtype of autism shares phenotypic characteristics with another disorder of the nervous system, we could repurpose treatments and accommodations working well in that other disorder and import them, adapting them to the autism subtype. In autism, we have a blanket treatment for all that is not working for many. This heterogeneous disorder with such homogeneous behavioral intervention has proven a poor model to aid neurodevelopment and is in fact stunting it. At the drug intervention level, we face a similar problem as we do with non-drug interventions. The broad heterogeneity that the current diagnostic criteria produce impedes tailoring interventions appropriately to the responsive features of the person’s nervous system. Subtyping autism into different categories, each one with similar genomic make up, could help us repurpose drug research in a more targeted manner (as explained in Figure 1.) For example, if we were to target genes responsive to certain compounds and those compounds were to alleviate symptoms of a physiological ailment, then knowing that those responsive genes are present in both a subtype of autism and another disorder (*e.g.*, ataxia) for which such compounds may have started research, we could leverage that research, bring it to autism, and advance drug discovery for a particular cluster sharing genes responsive to the compound. It is the same human brain and body for all, whether one has an autism label or not. Why not repurpose the scientific advances led by physiology and medicine in other fields, instead of being informed and guided exclusively by rather subjective psychological criteria? [33]

**Figure 1.**
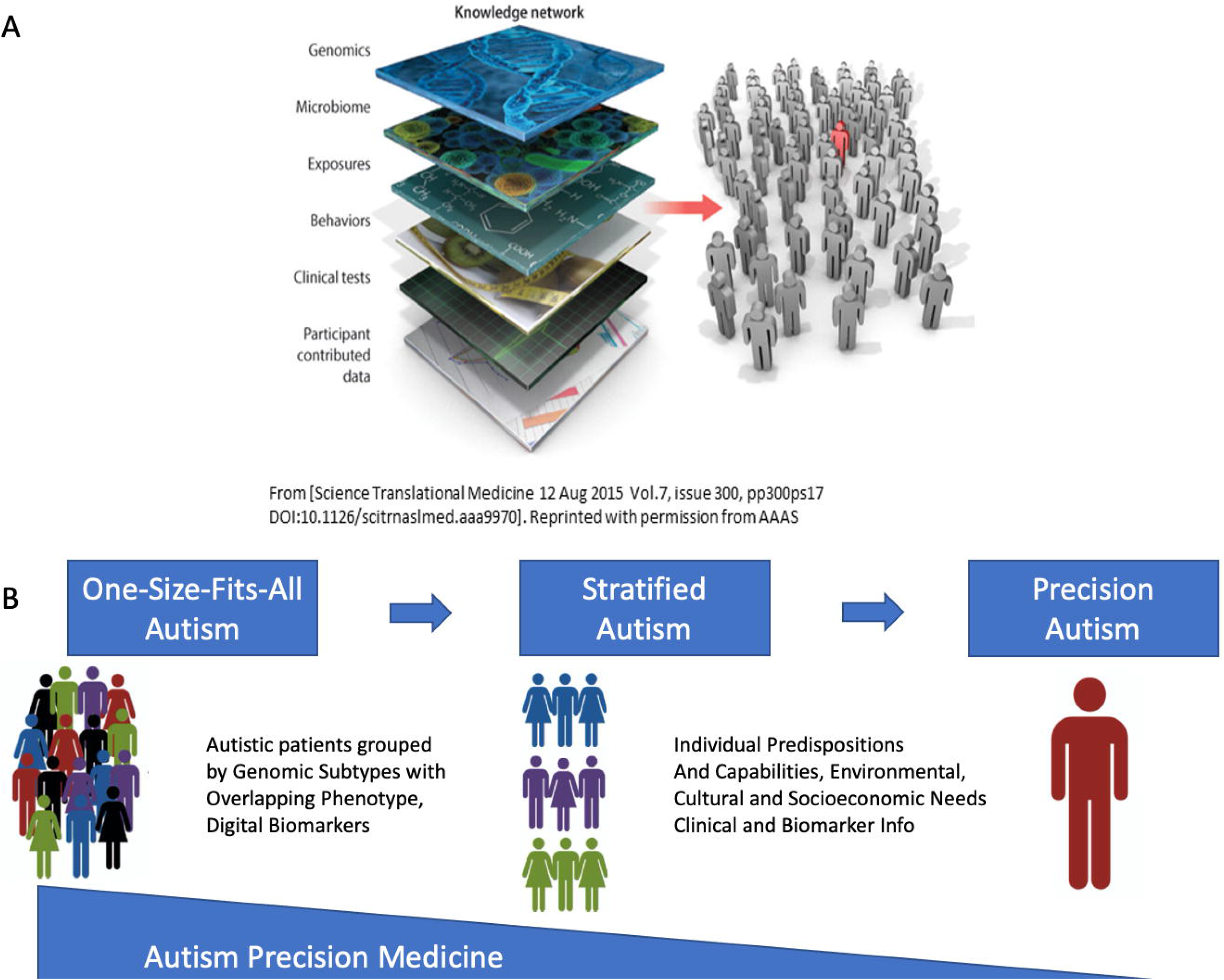
Leveraging existing genomic data to stratify the broad Autism Spectrum. (A) The Precision Medicine model [34] aims at the design of personalized targeted treatments that integrate all layers of the knowledge network to support the person’s needs under the genetic and epigenetic individual makeup. (B) Proposed model to stratify the broad spectrum of Autism based on existing genomic information causally defining the origins of neurological and neuropsychiatric disorders making up Autism today. This Precision Autism model can identify, relative to other disorders of known origin, the person’s best predispositions and capabilities, the environmental, cultural, and socio-economic needs, and design a personalized treatment that targets the medical-physiological issues rather than modifying behavior to conform to a grand average norm-a norm arbitrarily defined by current clinical criteria.

I find that we can indeed automatically and categorically stratify Autism through the gene pool of neurological and neuropsychiatric disorders that make up its broad spectrum today. I discuss our results in the context of the Precision Medicine (PM) model (Figure 1A) aimed at the development of personalized targeted treatments that integrate several layers of the knowledge network [34]. Under this PM platform I can better situate the person on the landscape of existing disorders for which there are more effective treatments than those currently offered in Autism. Based on this genomic stratification of Autism, I here propose a paradigm shift whereby the pipeline of diagnosis-to-treatment is based on the known physiology of these disorders (Figure 1B), addressing specific capabilities, predispositions and needs of each neurological phenotype. This new line of scientific inquiry not only leverages existing genomic information but more importantly, it responds to the quest of the autistic community to improve the prognosis and the future lives of those who receive this diagnosis.

## 2. Materials and Methods

I examine the genetic data base from the Simons Foundation Research Initiative (SFARI), which has been scored according to the evidence provided in the scientific literature linking the behavioral Autism diagnosis with a pool of genes. I then extract from that set those genes that overlap with the pool of genes linked to other disorders that are fundamentally defined by somatic-sensory-motor issues. Among them I use the genes associated with cerebral palsy (CP), Dystonia, Attention Deficit Hyperactivity Disorder (ADHD), obsessive compulsive disorder (OCD), Tourette’s, the Ataxias (autosomal dominant, recessive and X-linked) and Fragile X (FX). I also use the genes associated with Parkinson’s disease (PD) [35, 36], as symptoms of Parkinsonism abound in autistic adults after 40 years of age [30, 37].

While the SFARI genes circularly depend on the ADOS and the DSM behavioral Autism criteria (*i.e*., they were obtained precisely based on those clinical inventories describing *presumed* socially inappropriate behaviors), the latter genes come from the disease-association network (the DisGeNet portal) which did not rely on an Autism diagnosis. These individuals are likely to receive an Autism diagnosis at present, because of the shifts and broadening of the criteria [23, 24, 38–42]. However, they have their own clinical definition and known genetic origins [19, 43].

In Autism research, when the person has both diagnoses (ASD and a neurological or neuropsychiatric one) the former is coined idiopathic Autism, whereas the latter are called Autism of known etiology. Yet in a random draw of the population, we have identified clusters differentiating subtypes by relying on gait [6, 17, 44], voluntary reaches [4, 28, 45, 46] and involuntary head motions [5, 7, 30, 47, 48].

I blindly took these genes associated to other disorders of the nervous system (clinically and physiologically defined) and interrogated them in terms of their expression on brain and bodily tissues. I asked how much overlapping one would find (if any) with the SFARI Autism-linked genes (coined hereafter SFARI-Autism) defined by observation and descriptions of behaviors.

Upon this compilation of genes from several sources in the DisGeNet portal and the literature, I used the Genotype-Tissue Expression (GTEx) project involving human RNA-seq, expressed in Transcripts Per Million (TCM) to examine the genes’ expression across the 54 tissues sampled in their database [49] (Figure 2A). Using this atlas of genetic regulatory effects across human tissues, I compare across diseases, for those genes overlapping with the SFARI Autism-linked genes, the common tissues where the expression of these genes is maximal. To zoom into these overlapping genes expressed on tissues from the GTEx project (Figure 2A), I included the 11 distinct brain regions along with other 7 tissues representative of three fundamental muscle types: cardiac, smooth, and skeletal (Figure 2B), supporting the generation and maintenance of all physiological processes underlying all human functions and behaviors. Specifically, brain tissues included: the amygdala, the anterior cingulate cortex, the basal ganglia (caudate, putamen, nucleus accumbens), the brain cortex and the brain frontal cortex, the cerebellum and the cerebellar hemisphere, the hippocampus, the hypothalamus, and the substantia nigra.

**Figure 2.**
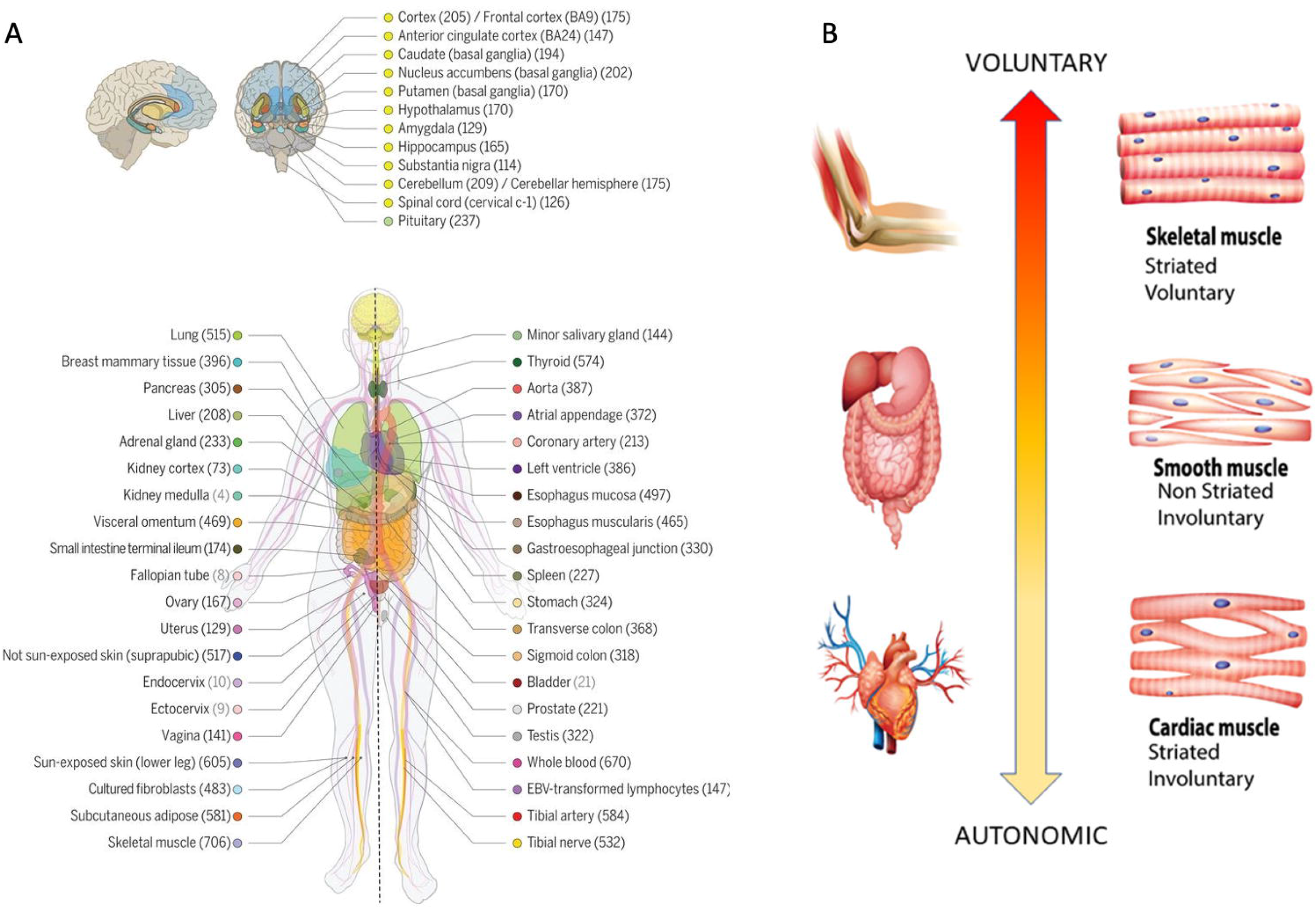
Genomic and Physiological criteria used in this study. (A) GTEx v8 study atlas of 54 tissues including 11 distinct brain regions and two cell lines. Genotyped sample donors numbers in parenthesis and color coding to indicate the tissue in the adjacent circles (Figure used with permission from AAAS [49].) (B) Three fundamental types of muscles supporting autonomic, involuntary and voluntary actions in humans can help us categorize behavioral functional levels according to related tissues affected [25].

Representative tissues of the different muscle types are: (cardiac) the heart left ventricle and the heart atrial appendage, (smooth) the esophagus muscularis and the bladder, (skeletal) muscle skeletal. Given their foundational role in all behavioral functions, I also examined these genes’ expression on the tibial nerve and the spinal cord (Figure 2A).

The SFARI Autism categories that I used to determine the level of confidence that the gene is linked to Autism, were those reported as of 03-04-2020. Quoting from their site:

### CATEGORY 1

Genes in this category are all found on the SPARK gene list. Each of these genes has been clearly implicated in Autism Spectrum Disorders, ASD—typically by the presence of at least three de novo likely-gene-disrupting mutations being reported in the literature—and such mutations identified in the sequencing of the SPARK cohort are typically returned to the participants. Some of these genes meet the most rigorous threshold of genome-wide significance; all at least meet a threshold false discovery rate of <0.1.

### CATEGORY 2

Genes with two reported de novo likely-gene-disrupting mutations.

A gene uniquely implicated by a genome-wide association study, either reaching genome-wide significance or, if not, consistently replicated and accompanied by evidence that the risk variant has a functional effect.

### CATEGORY 3

Genes with a single reported de novo likely-gene-disrupting mutation.

Evidence from a significant but unreplicated association study, or a series of rare inherited mutations for which there is not a rigorous statistical comparison with controls.

### SYNDROMIC (former category 4)

The syndromic category includes mutations that are associated with a substantial degree of increased risk and consistently linked to additional characteristics not required for an ASD diagnosis. If there is independent evidence implicating a gene in idiopathic ASD, it will be listed as “#S” (e.g., 2S, 3S). If there is no such independent evidence, the gene will be listed simply as “S”.

The GTEx dataset is as the 06-05-2017 v8 release [49]. For every gene in the disorders and diseases of interest, I first confirmed the presence of the gene in the GTEx dataset and then incorporated it into the analyses. This was necessary to provide the tissue expression from GTEx.

The genes from the DisGeNet portal were found by interrogation of their dataset under disease type and saving the outcome to excel files containing all pertinent information.

I follow our previously proposed roadmap to adapt the Precision Medicine platform [34] to Autism research and treatments [50] linking other disorders to the broad Autism phenotype (Figure 1). I first isolate the genes common to Autism and each disorder under consideration, sort them according to their median gene expression over the above mentioned 18 tissues of interest and then, for each tissue of interest, I highlight the top genes with expression above (log(e^60^) TMP (using the natural logarithm, Euler’s base 2.7183), to further help visualize common top genes across these diseases. I note that this is an arbitrary threshold used only to help visualize the top genes, since other thresholds could be used to visualize more genes in common expressing on tissues of interest. I report in the supplementary material the *full* set of genes common to these disorders and the SFARI-Autism set. Then, for each of the 54 tissues, I obtain the gene in the unique intersection set with the maximal expression and plot this information for the brain tissues of interest along with the SFARI-Autism score assigned to that gene.

In addition to genes linked to neurological disorders, I examined genes linked to neuropsychiatric conditions such as depression, schizophrenia, ADHD, and post-traumatic stress syndrome (PTSD), the latter owing to the tendencies in ASD to develop trauma and depression reportedly induced by current behavioral therapies [51], and to the known overlap between Autism and schizophrenia [52]. Furthermore, since the DSM 5 now accepts ASD and ADHD as coexisting diagnoses, I included ADHD-linked genes and asked about their overlap with Autism. I also tallied the shared genes pairwise across all the neurological and neuropsychiatric disorders under consideration, to learn about shared genes across these diseases.

Finally, other non-neurological and non-neuropsychiatric diseases were considered, to ascertain their overlap with the Autism-linked genes and with the genes linked to the neurological and neuropsychiatric disorders. I tallied their genes in common and interrogated the genes’ expression reported in the GTEx tissues. These included several forms of cancer (colon 49 genes, breast 488 genes, pancreas 114 genes), diabetes 5,545 genes, autoimmune disorders (lupus systemic 1,743 genes, psoriasis 1,221 genes, irritable bowel syndrome 1,483 genes), and congenital heart disease 252 genes, totaling 10,895 genes in addition to 10,028 genes associated to neural disorders (a random draw across 25 diseases of 20,923 genes and their expression on 54 tissues). Here I hypothesized that the overlap between the genes linked to Autism and those linked to neurological and neuropsychiatric conditions would be much higher than the overlap between the Autism-linked genes and the genes linked to other non-neurological diseases.

#### Methods to Obtain Pairwise Comparisons of Genes’ Expression in Autism and Various Disorders

I obtained for each set of reported DisGeNet genes liked to each disorder, their expression across the 54 tissues from the GTEx project. This yielded a matrix of N genes × 54 tissues, where each entry in the matrix is the gene’s expression in that tissue. Taking the median across all rows for each column (*i.e*., the number of genes in the disorder) gives a 1 × 54 vector array of median genes’ expression per tissue, which I normalize by the total number of genes in that disorder (scaling it to range between 0 and 1 unitless quantity). This is depicted in stem form in Figure 3A for each of 3 different representative disorders (ASD, ADHD and Lupus Systemic.) I then take the histogram of the values (represented by red dots in Figure 3A) and obtain the Earth Mover’s Distance (EMD) [53–55] between histograms, to code the distance in some probability space where I can represent these histograms according to an empirically fit continuous family of probability distributions. I obtain the EMD quantity pairwise between disorders, normalize it by the maximum value across the entire set, and represent it in matrix form for neuropsychiatric, neurological disorders and non-neurological diseases. I ask if clusters self-emerge from this representation of the median genes’ expression across the 54 tissues.

**Figure 3.**
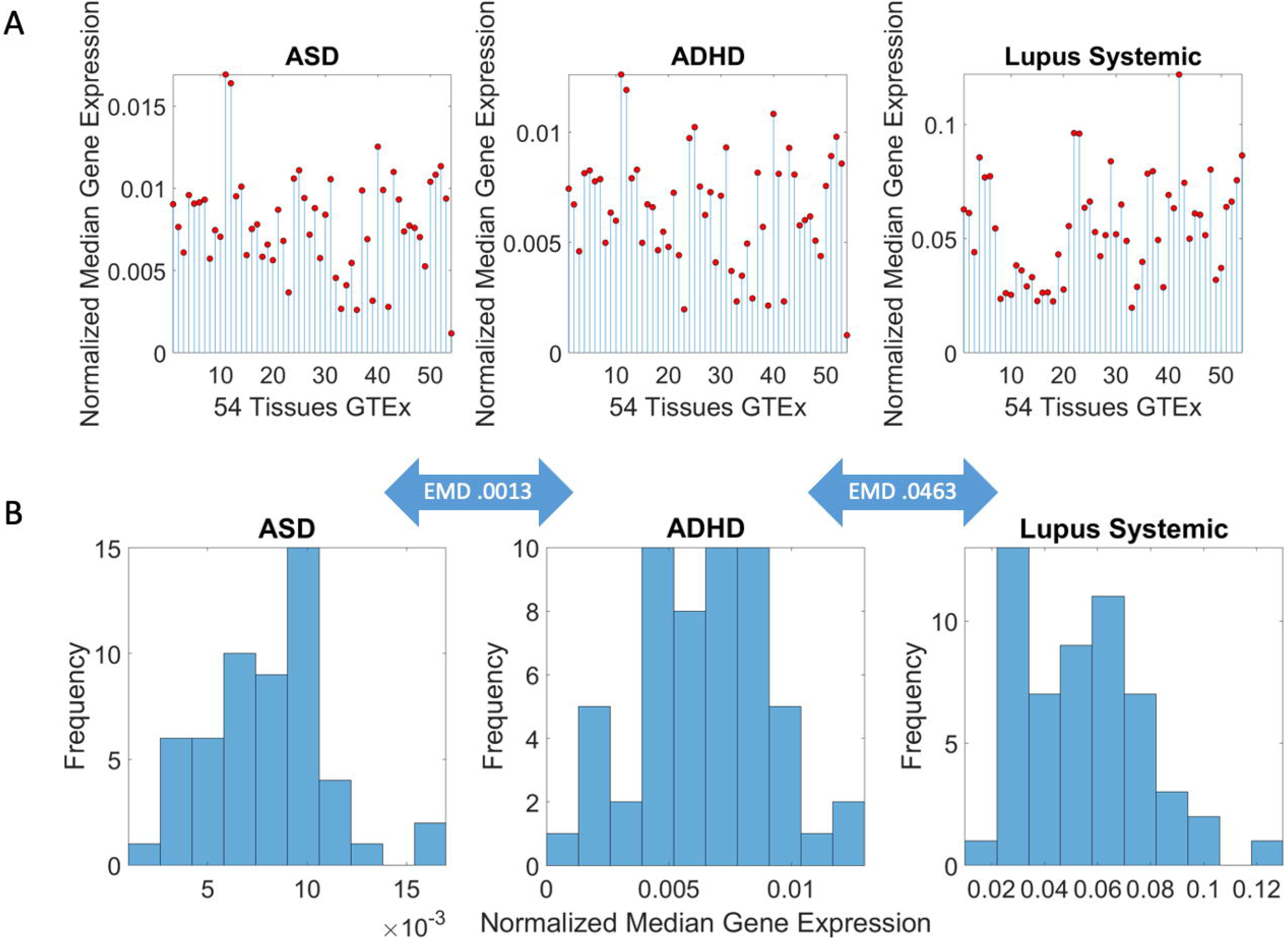
Pipe line of analysis to obtain pairwise similarity measurements between disorders. (A) The matrix of N genes × 54 tissues whereby is ij entry is the gene expression from the row i in the tissue j, is transformed into a 1 × 54 vector of median gene expression values (across the matrix rows) represented here in stem form, from the normalized values accounting for the number of genes in each disorder. Each red dot represents the median gene expression at the jth tissue. Three representative disorders are shown. (B) Histogram of the genes’ normalized expression correspond to the stem plots of panel (A). I take the earth mover’s distance (EMD) pairwise between two disorders (represented in the arrow) and build a matrix of EMD values representing the distance (similarity) between two disorders in the precise sense of genetic information contributing to genes’ expression on tissues.

## 3. Results

### 3.1. Neuropsychiatric Disorders and Autism Share Common Genes Expressed in Brain Tissues for Motor, Emotional and self-Regulatory Control

Quantification of genes common to Autism and neuropsychiatric disorders is depicted in Table 1. Figure 4A shows the pairwise shared genes color-coded (in log N color scale, where N is the number of genes in common with the Autism-linked SFARI set.) The inset in Figure 4A shows the distribution of genes common to the Autism linked SFARI database and each of the neuropsychiatric disorders under consideration, schizophrenia, ADHD, depression, and bipolar depression and including the neurological disorders. Notice that ADHD and Schizophrenia share the highest number of genes followed by depression and bipolar depression. Interestingly, I included PTSD owing to the tendency of trauma reported in Autism [51, 56, 57] and found 55 genes of those linked to PTSD in the SFARI-Autism set. I also included lupus systemic, owing to the known relations between autism and autoimmune disorders [58, 59].

**Table 1.** Overlap between Autism-linked genes in SFARI and known neuropsychiatric disorders, ranked by the % of genes obtained relative to the number of genes in the SFARI set under consideration here.

| Neuropsychiatric Condition | Number of Genes in DISGENET | Number of Genes Shared with SFARI ASD-linked Genes | % (Relative to DisGeNet, relative to 906 SFARI genes) |
| --- | --- | --- | --- |
| Schizophrenia | 2,697 | 336 | (24.58, 37.08) |
| ADHD | 795 | 188 | (23.64, 20.75) |
| Depression | 1,407 | 158 | (11.22, 17.43) |
| PTSD | 395 | 55 | (13.92, 6.07) |
| Bipolar Depression | 116 | 33 | (28.34, 3.6) |

**Figure 4.**
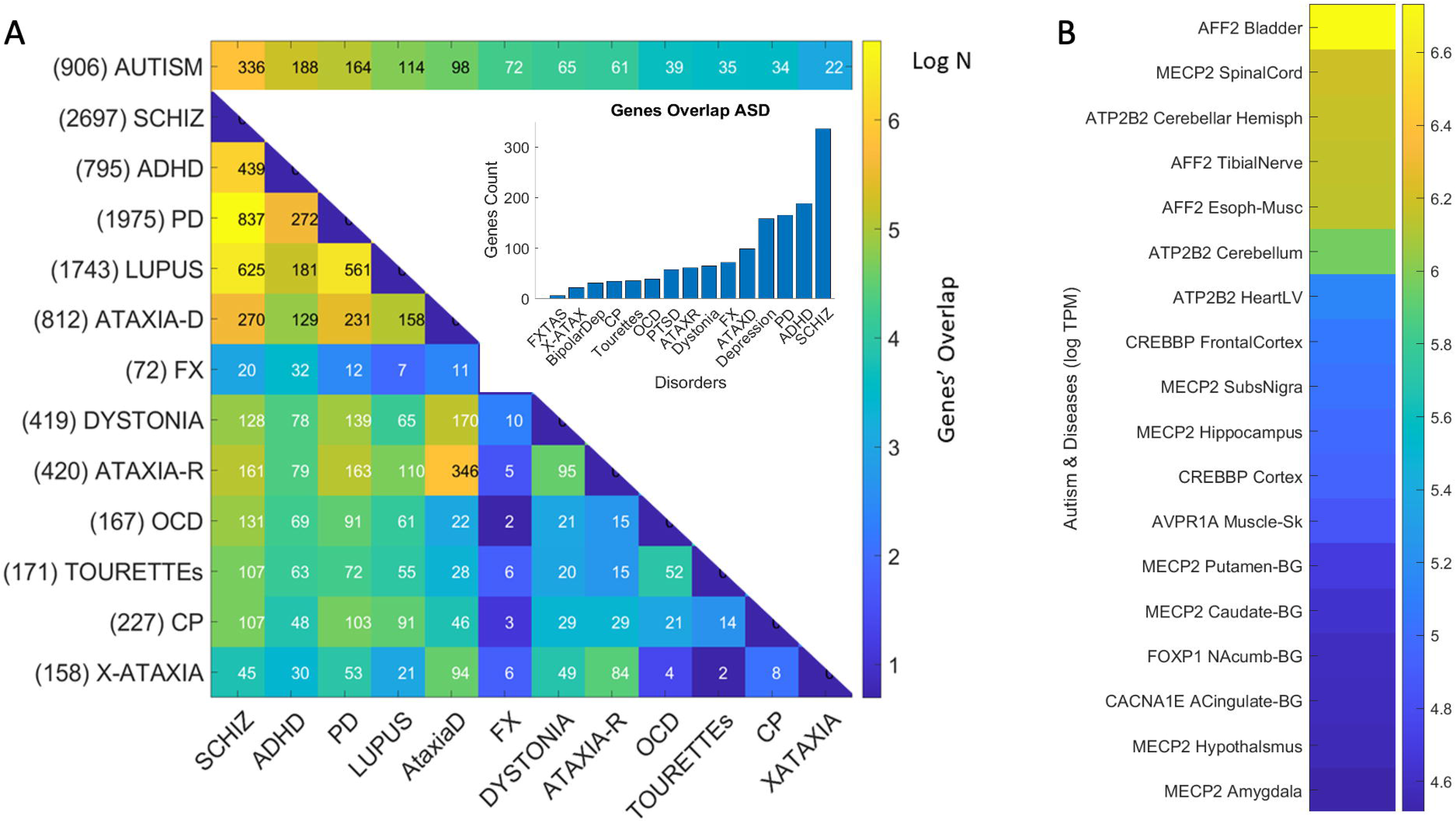
Number of genes common to Autism and selected neuropsychiatric, neurological, and autoimmune disorders. (A) Inset shows bar plot tallying the number of genes in each disorder that are also present in the SFARI dataset linking genes to Autism according to a confidence score (see methods.) Colormap depicting the number of genes in Autism and each disorder on the top row. Pairwise shared genes between disorder in row *i* and column *j* is the number on each entry of the matrix. Color is in log N, where N is the number of genes common to Autism and the disorder, or common to a disorder and another disorder (pairwise intersect.) (B) Color bar reflecting the 18 tissues with maximal gene expression and the corresponding gene at the intersection of SFARI-Autism and all shared genes in (A.)

### 3.2. Neurological Disorders and Autism Share Common Genes Expressed in Brain Tissues for Motor, Emotional and Regulatory Control

Likewise, quantification of genes linked to well-known neurological disorders and present in the SFARI-Autism dataset yielded up to 164 overlapping genes. These are depicted in Tables 1–2 for neuropsychiatric and neurological disorders respectively. I ranked them in each category by % relative to the 906 genes in the SFARI set under consideration, and by the number of genes associated to each disorder in DisGeNet. The shared genes are also shown in Figure 4A, color coded according to the number of genes in the intersection of SFARI-Autism and each disorder, and between disorders, taking the pairwise intersection. Table 1 shows that among the neuropsychiatric conditions, schizophrenia is the one with the highest percentage of genes shared with the SFARI-Autism set. Table 2 shows that among the neurological conditions, Parkinson’s disease has the highest percent shared with the SFARI-Autism set. This result came as a surprise, but it helps explain why as autistics age, the onset of Parkinson-like symptoms is reported by 40 years of age in 20% of the autistic adult population. This contrasts with .09% after 65 years of age in the general population [37]. Furthermore, this shared genetic pool between SFARI-Autism and the DisGenet genes associated to Parkinson’s disease helps explain the marked stochastic shift away from typical ranges of noise levels found in autistics at 40 years of age, when examining their involuntary head micro-motions at rest [30].

**Table 2.** Overlap between Autism-linked genes in SFARI and known neurological disorders of genetic origins ranked based on % relative to the set of 906 SFARI genes under consideration here.

| Neurological Disorder | Number of Genes Reported in DISGENET | Number of Genes Shared with SFARI ASD-linked Genes | % (Relative to DisGeNet, relative to 906 SFARI genes) |
| --- | --- | --- | --- |
| Parkinson's | 1,975 | 164 | (8.3, 18.10) |
| Ataxia Autosomal Dominant | 812 | 98 | (12.06, 10.81) |
| FX | 72 | 72 | (100.0, 7.94) |
| Dystonia | 419 | 65 | (15.51, 7.17) |
| Ataxia Autosomal Recessive | 420 | 61 | (14.52, 6.73) |
| OCD | 167 | 39 | (23.35, 4.30) |
| Tourette's | 171 | 35 | (20.46, 3.86) |
| CP | 227 | 34 | (14.97, 3.75) |
| X-Ataxia | 158 | 22 | (13.92, 2.42) |
| FXTAS | 63 | 22 | (34.9, 2.42) |
| Progressive Cerebellar Ataxia | 134 | 13 | (9.70, 1.43) |

Surprisingly also was the finding concerning schizophrenia in Figure 4A, whereby 837 genes are shared between Parkinson’s disease and the schizophrenia set, while 439 are shared between ADHD and schizophrenia. Furthermore, 625 genes are shared between lupus systemic and schizophrenia. This figure depicts the shared gene pool across these selected disorders that also show their genes overlapping with Autism-linked genes (on the top row.) The numbers next to the disorder are the number of genes reported in DisGeNet. The numbers in the color map entries are those shared pairwise between the disorders in row i and column j of the matrix. Figure 4B shows the tissues with the maximal gene expression using a color bar (in log median TCM) sorted in descending order. These are the genes common to all the disorders under consideration that overlap with SFARI-Autism genes.

### 3.3. Examination of the Maximal Gene Expression on the Tissues for Genes Common to Autism as Revealed a Compact Gene Pool

I found that 12 genes common to Autism and these disorders are maximally expressed on the 54 tissues. They are depicted in Table 3 along with the tissues where they maximally express, while Figure 4B shows the genes maximally expressed on the 18 tissues of interest (brain, spinal cord, tibial nerve, and those key to cardiac, smooth, and skeletal muscles.) I discuss later some of the literature on these known genes. I also provide supplementary text files containing for each disorder, the pairwise genes in common between Autism and each disorder or disease under consideration.

**Table 3.** Compact set of genes common to Autism and the neurological and neuropsychiatric disorders maximally and selectively expressed across the 54 tissues.

| Genes Common to ASD and Neuro-Disorders | Tissue with Max Expression |
| --- | --- |
| ACTB | Liver |
| AFF2 | Adipose Subcutaneous, Adipose Visceral Omentum, AdrenalGland, Artery Aorta, Artery Coronary, Artery Tibial, Bladder, Breast Mam Tiss, Cervix Ectocervix, Cervix Endocervix, Colon Sigmoid, Colon Transverse, Esophagus Gastro esophageal Junction, Esophagus Muscularis, Fallopian Tube, Heart AtrialAppendage, Kidney Cortex, Kidney Medulla, Lung, Minor Salivary Gland, Nerve Tibial, Pituitary, Prostate, Small Int ileum, Spleen, Stomach, Thyroid, Uterus, Vagina |
| AKAP9 | Whole Blood |
| ALDH5A1 | Esophagus Mucosa |
| ATP2B2 | Heart Left, Ventricle, Ovary, Cerebellar Hemi, Cerebellum |
| AVPR1A | Muscle Skeletal |
| CACNA1E | Ant Cingulate Cortex |
| CHD7 | Pancreas, Skin not Sun Exposed Suprapubic, Skin Sun Exposed Lower Leg |
| CREBBP | Cortex, Frontal Cortex, Fibroblasts, Cell-Lymphocytes |
| FOXP1 | Nucleus Accumbens of the Basal Ganglia (BG) |
| MECP2<br>G | Amygdala, Caudate-BG, Hippocampus, Hypothalamus, Putamen-BG, Spinal Cord-Cervical, Substantia Nigra |
| SMAD4 | Testis |

### 3.4. Genomic Stratification of Neurological and Neuropsychiatric Diseases Making Up Autism Today

The EMD taken pairwise between Autism and each disorder, and pairwise acrossall neuropsychiatric, neurological disorders and non-neurological diseases revealed an orderly stratification of disorders, whereby a common gene pool and expression on the tissues can clearly separate neuropsychiatric and neurological from non-neurological diseases. This is shown in Figure 5A, where we can also appreciate that in the non-neurological diseases, the autoimmune ones share a common gene pool and tissue expression. Notably, colon cancer is also close in a probability distance sense, to neurological disorders of known genetic origin, namely, Fragile X, FXTAS, the ataxias, dystonia, and Parkinson’s disease.

**Figure 5.**
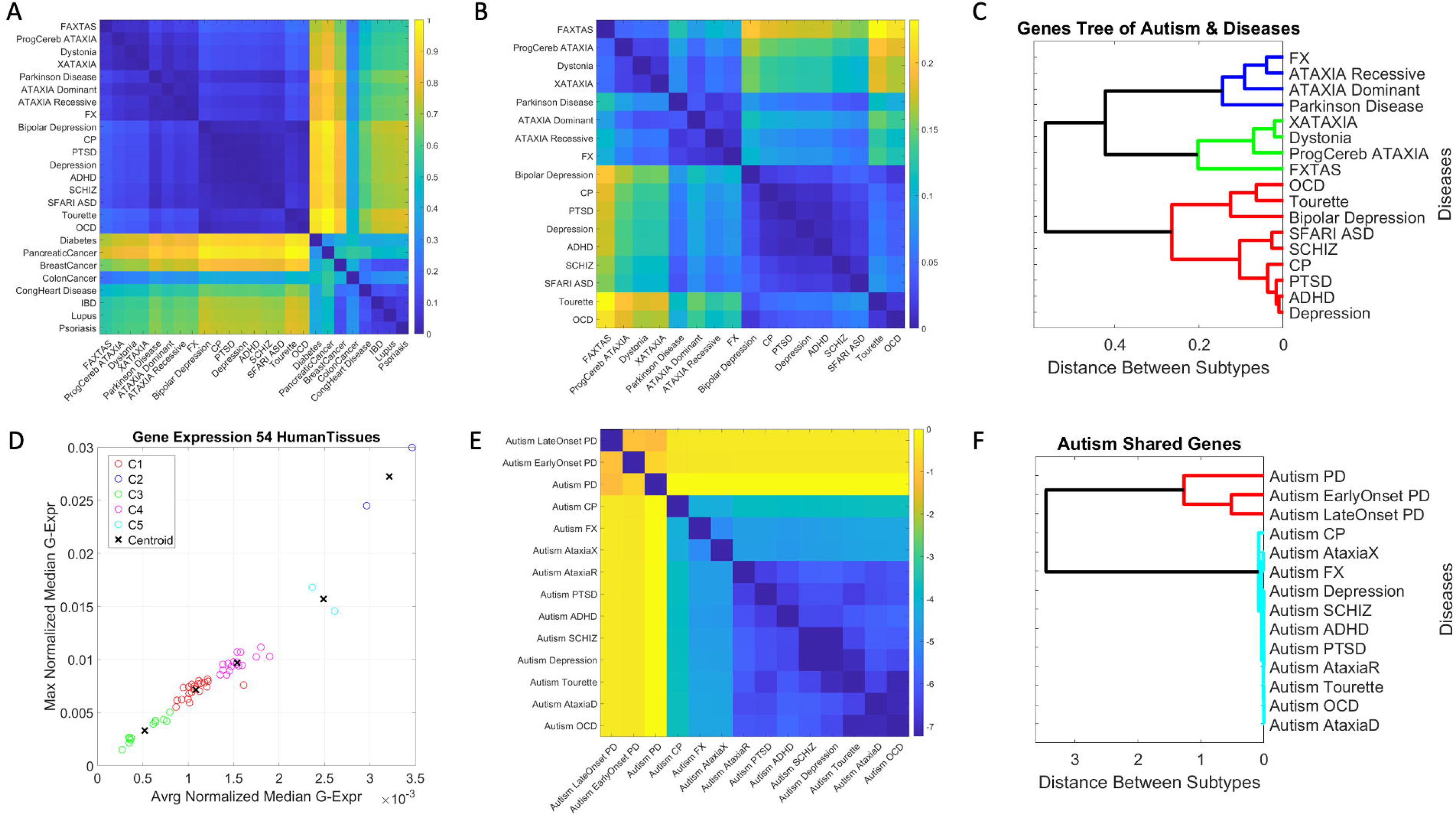
Genomic stratification of Autism and diseases obtained by leveraging the gene pool that overlaps with other known neurological and neuropsychiatric diseases and their expression on the 54 tissues defined by GTEx. (A) EMD-based separation between neuropsychiatric, neurological disorders and non-neurological diseases, also identifying common gene pool in autoimmune diseases. Color scale is the normalized EMD value taken pairwise between the vector of median gene expression across 54 tissues of GTEx. (B) Zooming into the neuropsychiatric and neurological diseases whose gene pool overlaps with Autism, I see different self-emerging subclusters further refining the stratification. (C) Dendogram showing the binary tree orderly organization that groups and categorizes diseases according to genes’ overlap and tissue expression with respect to ASD. (D) Output of K-Means algorithm with 5 tissue-cluster criteria taken on shared genes between SFARI-Autism and each disorder/disease in (B-C). Cluster 1 (red) includes the amygdala, hippocampus, putamen, and substantia nigra. Cluster 2 (blue) in a category of its own, includes the cerebellar hemisphere and the cerebellum. Cluster 3 (green) does not contain brain tissues but contains tissues important for cardiac (heart atrial appendage, heart left ventricle), smooth (esophagus mucosa, bladder) and skeletal muscles (muscle skeletal.) Cluster 4 (magenta) contains the anterior cingulate cortex, the basal ganglia’s caudate and nucleus accumbens, the brain cortex, the hypothalamus. Cluster 5 (cyan) contains the frontal cortex and the pituitary gland. (E) Similarity matrix built by taking the normalized Earth Mover’s distance metric pairwise between the genes in the intersection of Autism and each of the 14 disorders under consideration. Higher distances (yellow) indicate more effort to transform the distribution of median values of genes’ expression (taken across the 54 tissues) from one set of SFARI-Autism shared genes and a given disorder, with another set of SFARI-Autism shared genes and another disorder. (F) Hierarchical clustering of these shared genes identifies two main groups of shared genes with SFARI-Autism, one formed by those SFARI-Autism genes found in the genes linked to PD, early onset PD and late onset PD, and the other formed by the genes shared (pairwise) with each of the other neuropsychiatric and neurological disorders under consideration.

Zooming into the entries with lowest EMD value, corresponding to the neuropsychiatric and neurological disorders, we see in Figure 5B, that other patterns self-emerge further refining the clusters. There, we can appreciate that ASD is close to ADHD and Schizophrenia, as well as close to Depression, PTSD and Cerebral Palsy. Furthermore, OCD and Tourette’s cluster close together, also showing a common gene pool and genes’ expression across the tissues. In the group of the neurological disorders of known etiology, we can visualize self-grouping of FX and the ataxias (dominant and recessive), while X-ataxia, dystonia, Progressive cerebellar ataxia, and Parkinson self-cluster and separate from FXTAS.

To further test this visualization, I ran a common clustering procedure using MATLAB linkage function applying Euclidean distance and plotting the output as a dendogram. This shows the orderly binary tree structure of these genes-tissues grouping in Figure 5C. We can see that there are three main subtrees of the binary tree, *i.e*., two subtrees comprised of neurological disorders, and one with neuropsychiatric disorders of the types diagnosed by the DSM. Further refinement reveals FXTAS as a leaf of its own, close to progressive cerebellar ataxia, dystonia, and X-ataxia, all under the same subtree. The other subtree contains Parkinson’s disease, ataxia dominant and recessive and Fragile X.

At the neuropsychiatric end, we see that Tourette’s and OCD group in a branch and bipolar depression is a leaf of its own, while schizophrenia and SFARI-Autism fall the closest together in one branch of the same subtree. That subtree also groups PTSD, Depression and ADHD under one branch and shows CP as a leaf of its own. These gene pools and their expression on the 54 GTEx tissues define an orderly stratification aided by genes common to autism spectrum disorders, according to DisGeNet and SFARI genomic reports.

Clustering by tissues maximally expressing the shared genes across disorders (Figure 5D), we can see 5 groups of tissues whereby, the cerebellum and cerebellar hemisphere are by far the tissues with the highest gene expression, followed by the prefrontal cortex and pituitary gland. The following group is comprised of the anterior cingulate cortex, the basal ganglia’s caudate and nucleus accumbens, the brain cortex, the hypothalamus, followed by the cluster containing the amygdala, hippocampus, putamen, and substantia nigra. The lowest expression is in the cluster containing no brain tissues, but tissues important for survival and overall functioning of cardiac (heart atrial appendage, heart left ventricle), smooth (esophagus mucosa, bladder) and skeletal muscles (muscle skeletal.)

Furthermore, I examined the pairwise intersection sets of genes shared between SFARI-Autism and each of these disorders in the neurological and neuropsychiatric sets. These are found in Figure 5E, as they grouped according to the EMD metric (expressed here in log scale.) The hierarchical clustering revealed two main subtrees in this case, one comprising PD and early and late onset PD in the intersection with SFARI-Autism. The other branch revealed an orderly grouping of neurological disorders surrounding the neuropsychiatric disorders (depression, schizophrenia, ADHD, and PTSD.)

Figures 6 and 7 reveal the genes expression for values above (log(e^60^) TCM) with the SFARI confidence score on the horizontal axis and the expression value log TCM on the vertical axis. Figure 6 shows the brain tissues and genes above this level of expression along with the confidence score in the SFARI data repository. Tissues that are fundamental for the development and maintenance of motor learning, motor coordination and motor adaptation include the substantia nigra (maximal expressed gene AFF2-AUT1 signifying this gene is scored in SFARI as score 1 confidence), basal ganglia with caudate, putamen and nucleus accumbens also with AFF2-AUT1 as top expressing gene. Other tissues involved in motor control are the cerebellar hemisphere (ATP2B2-AUT2) and the cerebellum (CREBBP-AUT1). Tissues known to be involved in executive function are the brain frontal cortex (MECP2-AUT1) and the brain cortex (CACNA1E-AUT2). Tissues known for their involvement in emotions (amygdala (AFF2-AUT1)) and memory (hippocampus (BRSK2-AUT1)) and the anterior cingulate cortex (BRSK2-AUT1) are also depicted in Figure 6, along with the schematics of the brain from Figure 1 with the locations of these areas.

**Figure 6.**
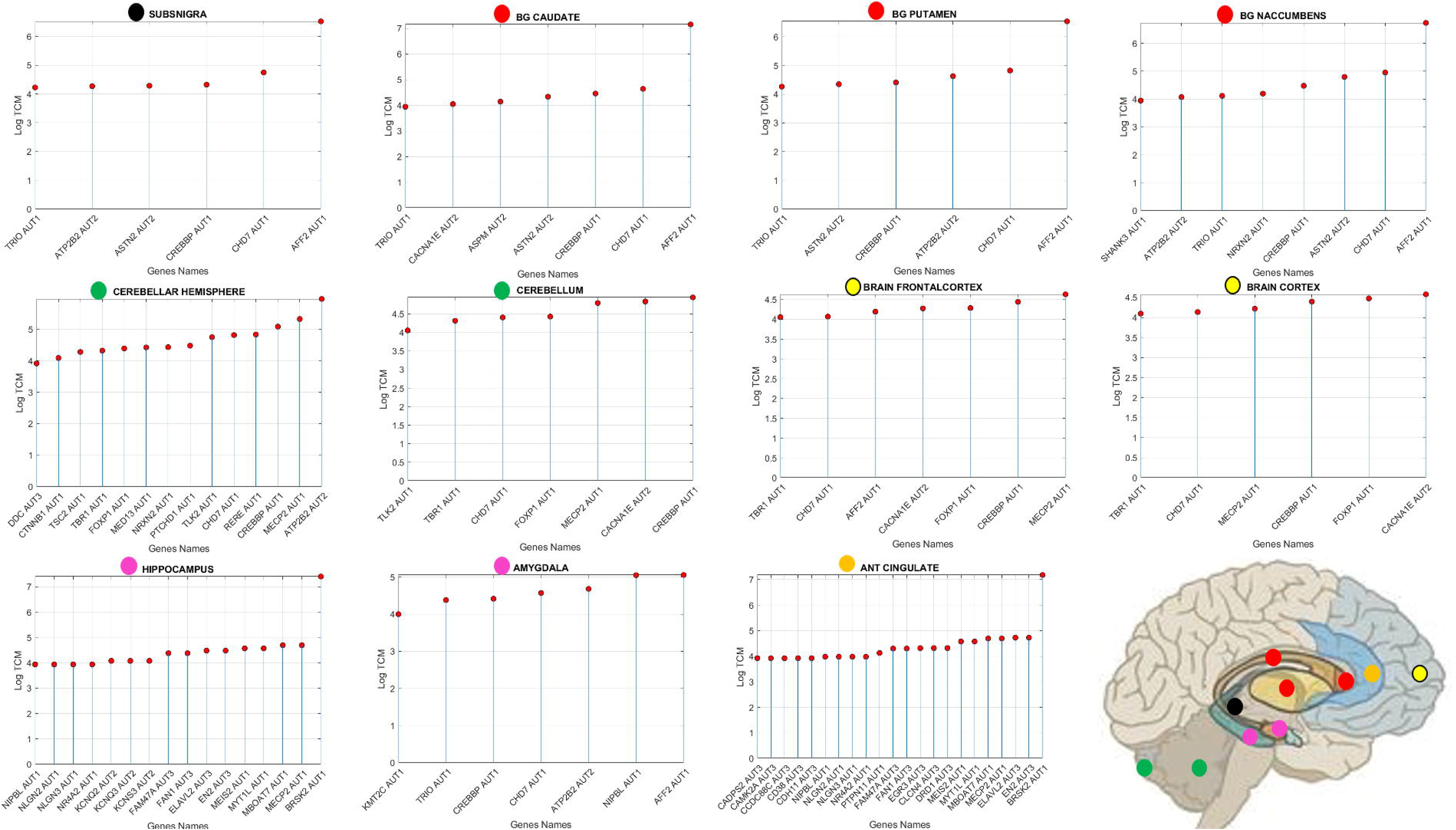
Common genes to Autism and all other neurological and neuropsychiatric disorders. maximally express on brain tissues involved in the initiation, generation, control, coordination, and adaptation of movements, as well as in tissues necessary for the creation, retrieval and maintenance of memories and executive function. Genes’ expression is threshold above log(e^60^) to show the top expressing genes from DisGenNet overlapping with those in the SFARI set (Full list of shared genes are in the Supplementary material files.) The AUT# reflects the confidence score assigned to the gene at the SFARI repository. Horizontal axis shows the genes above threshold and vertical axis gives the expression level (log transcripts count per million, TCM.) Each colored dot is shown at the brain tissue in Figure 2A schematic form.

**Figure 7.**
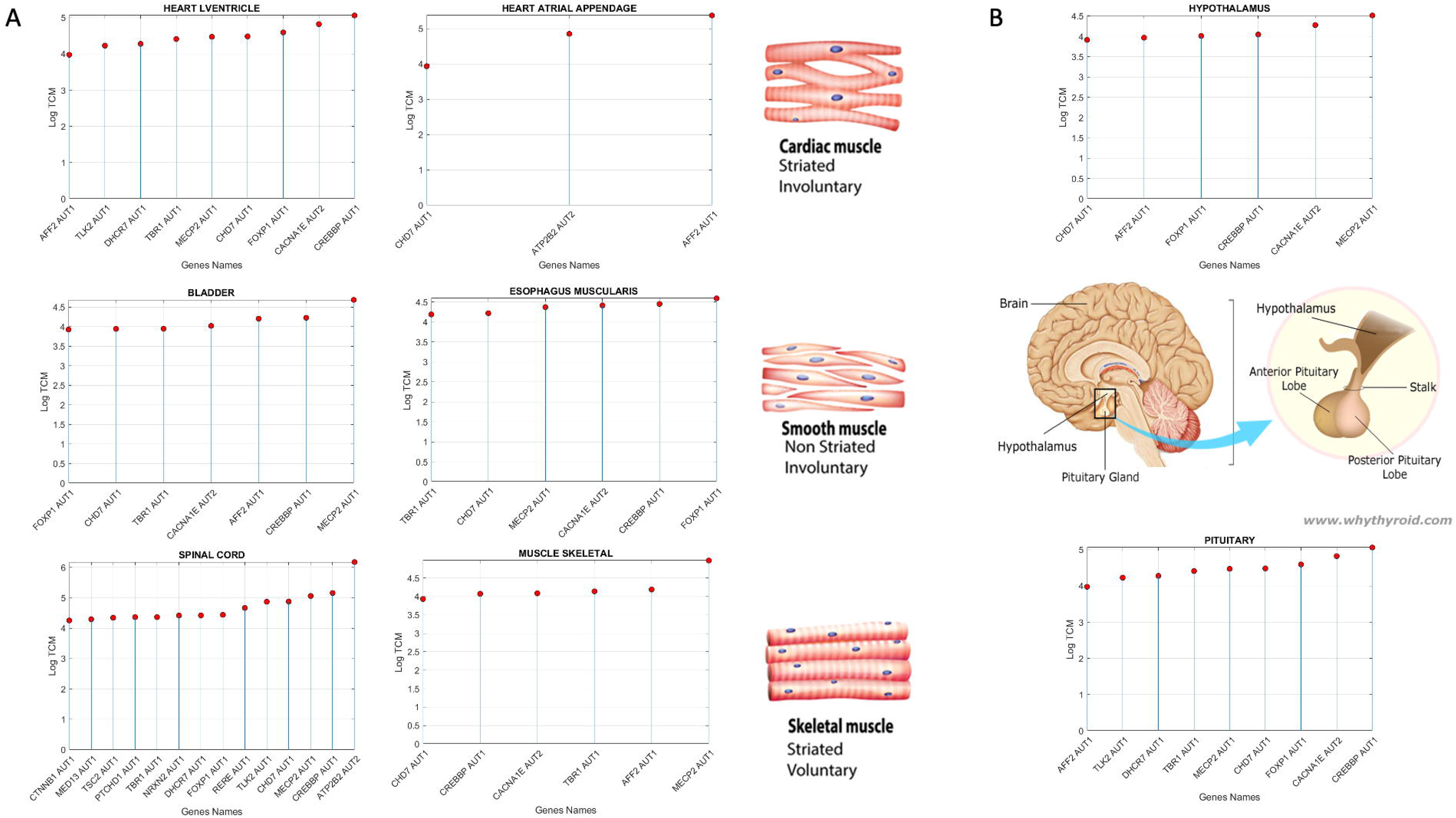
Genes present in Autism and all other neurological and neuropsychiatric disorders. maximally expressed (above log(e^60^) on tissues involved in all vital functions associated with cardiac, smooth, and skeletal muscles and the spinal cord (A), and with self-systemic regulation (B). Top genes and corresponding SFARI confidence score are also plotted on the horizontal axis. Vertical axis reflects the expression in log TCM units. Schematics reflect the locations of the hypothalamus and pituitary gland in the brain (from www.whythyroide.com)

In Figure 7, I also reveal those genes’ expression and SFARI scores for the cardiac (heart left ventricle and the heart atrial appendage), skeletal (muscle skeletal), and smooth muscle (esophagus muscularis and bladder) tissues and for the nerve tibial. The latter is critical to develop kinesthetic reafference and proper gait, known to be disrupted in several of these disorders (autism, PD, FX [44]). Common to all these disorders are well known genes in the Autism literature with SFARI score confidence 1 (*e.g*., MECP2, AFF2, FOXP1, CREBBP, CACNA1E, CHD7, TRIO and SHANK3, among others.) I will later discuss the known roles of some of these genes in the development of synapses and circuits necessary to form and dynamically maintain neural networks.

I further plot in Supplementary Figures 1-3 genes common to Autism and some of the disorders, for the top 20 genes with maximal expression (in log TCM units) across the brain and spinal cord tissues, as well as tissues involving muscle skeletal, cardiac, and smooth muscle types. These figures in the Supplementary Material show the matrix with genes across the rows (top 20 expressed genes) and 18/54 tissues across the columns. Each entry is color coding the expression of these genes in log TCM units. From these plots, I note that e.g., MECP2 is present at the highest expression on the spinal cord in Autism and Schizophrenia, Autism and Depression, Autism and ADHD, Autism and OCD, Autism and Cerebral Palsy, Autism and Dystonia, Autism and Autosomal Dominant Ataxia, Autism and Autosomal Recessive Ataxia, Autism and Fragile X, Autism and X-ataxia, Autism and PTSD but not in Tourette’s & Autism, where MECP2 is not among the top 20 genes expressed on these tissues. Instead, CHD2 is expressed on the spinal cord, and highly expressed on the cerebellum and cerebellar hemisphere. Indeed, Tourette’s is closer to OCD (Figure 5C) than to the cluster formed by ASD, ADHD and Schizophrenia (though located on the same subtree as these neuropsychiatric disorders, but in a separate branch containing bipolar depression too.)

## 4. Discussion

This work provides support to the idea of reframing Autism under the model of Precision Medicine [25, 50], while also addressing the notion of an “Autism epidemic” recently portrayed as an exponential rise in prevalence [60] and its costly consequences [60]. Re-examining Autism as the conglomerate of disorders and diseases, many of known origins, comprising this heterogeneous spectrum, I conclude that such “*epidemic*” or “*tsunami*” is bound to be an artifact of the current behavioral diagnosis-to-treatment pipeline. This pipeline allows such comorbidities and often shifts criteria, discouraging stratification. I invite the thought that stratifying the spectrum according to underlying genetic, causal information would provide far more viable strategies to cope with the overall increase in neurodevelopmental disorders in general, than continuing the use of Autism as a blanket label grouping all these disorders. Furthermore, I argue that several of the disorders in question already have therapies designed to address issues in their corresponding phenotype. As such, the general Autism label, when funneled through genetic subtypes, could leverage the accommodations, and offer support pertinent to each of the neurological and neuropsychiatric groups conforming its broad spectrum today. Here we report that they have a considerable genetic overlap with the genes linked to SFARI-Autism.

Autism serves as an umbrella term encompassing many disorders and diseases, some of which have precise etiology (*e.g.*, Down Syndrome [61], Fragile X Syndrome [62], *etc*.) I therefore combined multiple open access data sets with the label of Autism and with the label of a disorder or disease that often receives the Autism diagnosis. I included neuropsychiatric disorders, and neurological and non-neurological diseases associated with some sets of genes. Then, I applied common computational techniques to attempt to automatically and orderly stratify a cohort of 25 diseases and 20,923 genes expressed on 54 brain and bodily tissues, vital for the survival and functioning of the individual

I show that given a random draw of genes linked to disorders with high penetrance in Autism, and even some which are not officially associated with autism, one could find self-emerging clusters at their intersection. Using (probabilistic) distance metric assessing the similarity between genes associated to autism and those associated to the other disorders, and examining their expression in brain-body tissues, several important patterns self-revealed. Among them, DisGeNet Parkinson’s disease emerged as the neurological condition with maximal number of shared genes associated to the SFARI-Autism set under consideration. Schizophrenia appeared as the neuropsychiatric disorder with maximal number of genes shared with Parkinson’s disease. Both SFARI-Autism and Schizophrenia shared the maximum number of genes with the SFARI set, strongly suggesting that movement disorders are at the core of both autism and schizophrenia.

I found self-emerging clusters that clearly (and automatically) differentiated neuropsychiatric and neurological disorders from non-neurological diseases and within the brain-related disorders, I established an orderly distance to Autism in the sense of overlapping genes and their expression on tissues critical for motor control (initiation-termination, learning, coordination, sequencing, and adaptation), cognition, memory, and self-regulation. I also found that the autoimmune disorder lupus systemic shares 114 genes with those linked to Autism in SFARI (12.6% relative to the 906 genes in SFARI), a result congruent with recent links between Autism and autoimmune disorders [58, 59, 63].

The overall conclusions from these results are several folds: (1) Autism is a movement disorder and should be accordingly redefined and treated as such, rather than treated as a misbehavior; (2) The broad spectrum of Autism, as we know it today, *i.e*., inclusive of disorders and diseases of known etiology, share a common set of genes with genetic disorders and consequentially can be stratified into Autism subtypes. (3) Given this automatic clustering, it is safe to conclude that Autism prevalence rates are an artifact of current surveillance methods relying exclusively on the clinical (behavioral) diagnosis that welcomes other disorders. Incidentally, it has been shown that such methods of diagnoses use fundamentally flawed statistics in the criteria, thus inflating false positives [9]. Furthermore, digitizing them with wearable biosensors, captures the fundamental differences in females, saves time, thus being less taxing on the children and offering a new level of finer granularity of physiological function, well beyond the limits of the naked eye. As such, digitized dyadic interactions during the ADOS opens a new avenue for precision (physiological) phenotyping that, when combined with genomics results here, stands to reformulate autism research under the tenets of Precision Medicine [25, 64].

The genetically-based subtypes reported here might be more manageable and less costly to treat and service, than forecasted by current epidemiological accounts relying only on the psychological construct of “*appropriate*” social behaviors [60]. Since there are therapies for other disorders that share genes associated with Autism, it may be possible to repurpose such therapies and adapt them to corresponding Autism subtypes.

### 4.1. The Genetically Informed Autism Subtypes

Given a mixture of genes and their median expression on the 54 tissues defined by GTEx, across multiple neuropsychiatric and neurological disorders, I found that Autism-linked genes in SFARI overlap with genes defining those disorders in DisGeNet, which would also likely receive the Autism label. Such overlapping showed an orderly stratification using a binary tree structure that rendered schizophrenia as the closest to SFARI-Autism (sharing 363/906 genes reported in SFARI) and FXTAS as the farthest (nonetheless sharing 22/906 genes reported in SFARI.) I note that DisGeNet ASD (autism spectrum disorders; CUI: C1510586) is a superset of SFARI, 1,071 *vs*. 906 genes and SFARI genes are ranked according to the literature. I also note that since our last download, the number of genes in SFARI may have increased.

In good concordance with clinical criteria, two main subtypes automatically self-emerged according to the distance metric that I used (Figure 5C) comprising neuropsychiatric and neurological disorders, all sharing a compact set of genes described below. The neuropsychiatric branch includes the Autism linked genes in DisGeNet (overlapping with a subset of those in SFARI) and disorders in the Diagnostic Statistical Manual, 5^th^ edition, DSM-5 and the International Classification of Diseases, 10^th^ edition, ICD-10. In order of distance to SFARI-Autism, on one end we have schizophrenia (the closest sharing the same sub-branch), CP, PTSD, ADHD (allowed to be co-diagnosed with Autism in the DSM5) and depression. Then, the other branch has bipolar depression, Tourette’s, and OCD. Among the neurological disorders conforming the second cluster, I have in order of distance from the SFARI-Autism set, FXTAS, progressive cerebellar ataxia, dystonia, and X-ataxia. The last cluster has Parkinson’s disease, ataxia dominant, ataxia recessive and Fragile X, with Parkinson’s disease sharing the largest percent of SFARI-Autism genes. This natural breakdown of the (Autism) spectrum according to the shared genetic pool and genes’ expression on tissues fundamental to form the building blocks of any human behavior, is far more manageable (using physiological and medical knowledge today) than considering the full spectrum in a monolithic form, as suggested by psychological surveillance methods [1, 60] and the behaviorists’ recommendations [13].

### 4.2. The Genes Common to Autism and Each Subtype

Each neuropsychiatric or neurological subtype identified by the genes-tissue analysis shared genes with the SFARI-Autism set. The full list in Supplementary Materials for each disorder/disease, offers clues with regards to the tissues whereby these intersecting genes maximally express. Furthermore, Figure 5D revealed several clusters of tissues common to all these pairwise-shared genes between the disorders and SFARI-Autism. Among these clusters, the cerebellar hemisphere and the cerebellum emerged as a separate group, common to all these disorders, with the maximum average median gene expression. This cluster of tissues was followed by a cluster that included the frontal cortex and the pituitary gland, also far apar from the other three clusters in Figure 5D.

Autism and schizophrenia, Autism and ADHD, Autism and Depression, Autism and OCD, Autism and FX, Autism and ataxia-X, Autism and PTSD, Autism and CP, Autism and dystonia, Autism and Ataxia autosomal dominant and Autism and Ataxia autosomal recessive, all share METHYL-CpG-binding protein 2, MECP2, with cytogenetic location at Xq28 (according to the Online Mendelian Inheritance in Man, OMIM site). It is reported as implicated in severe neonatal encephalopathy, mental disability and Rett syndrome, as well as to have high Autism susceptibility. MECP2, binds methylated CpGs. It is a chromatin-associated protein that can both activate and repress transcription; it is required for maturation of neurons and is developmentally regulated [65].

Furthermore, Autism and schizophrenia and Autism and ADHD (the two top neuropsychiatric disorders sharing the highest percentages of genes with the genes linked to SFARI-Autism) shared the CREP-Binding Protein (CREBBP), among the top 10 genes expressed maximally across the 54 tissues and common to the Autism and Schizophrenia gene pool. It is located in 16p12.3, a chromosomal region linked to Autism.

Autism and Tourette’s syndrome did not share MECP2, but shared CHD2 as the top gene maximally expressed across all 18 tissues of the brain, spinal cord and tissues associated with cardiac, smooth, and skeletal muscles. Maximal expression at the cerebellar hemisphere and the cerebellum suggests involvement in motor control, coordination, and adaptation, while high expression in other tissues for memory, cognition, and self-systemic regulation suggest that this gene is rather important. Indeed, prior work in Autism [66] and other neurodevelopmental disorders [67] had conferred importance to this gene for neural development.

Among highly expressed genes in Autism and other disorders, I also found AFF2 [68] and BRSK2 [69], both with score 1 in SFARI and reportedly critical for neurodevelopment. Indeed, the X-linked gene AFF2 has been found in patients with fragile X E (FRAXE) intellectual disability, while the gene encoding the serine/threonine-protein kinase BRSK2 was recently detected in individuals with developmental and intellectual disability.

To further understand the possible links that have been suggested between Autism and PD (particularly during adulthood), I also examined the genes from DisGeNet linked to early (50 genes) and late (238 genes) onset of PD, along with those linked to PD in general (1,975 genes.) This revealed that DisGeNet PD shares 164 genes with SFARI-Autism, whereas early onset PD shares 8 and late onset PD shares 32 genes with those in SFARI-Autism. Among the genes maximally expressed in the tissues of the brain and the cardiac, smooth, and skeletal muscles in PD, AFF2 and TSC2 were found. In early onset PD, RAB39B and SLC6A3 were found. Mutations in RAB39B cause X-linked intellectual disability and early-onset Parkinson disease with alpha-synuclein pathology, also linked to X-linked mental disability associated with Autism, epilepsy, and macrocephaly [70–73]. SLC6A3 provides instructions for making the dopamine transporter protein (DAT) embedded in dopaminergic neurons. Variations (polymorphisms) of the SLC6A3 gene have been linked to PD, ADHD [74] and ASD [75]. Dopamine is a known neurotransmitter important for multiple cognitive and motor functions, as well as for the functioning of the reward systems of the brain. In late onset PD, TET2, ADA and PTGS2 (COX2) were found. Located in 4q24, TET methylcytosine dioxygenase 2 is listed in OMIM as a TET protein playing a key role in the regulation of DNA-methylation status serving both as a stable epigenetic mark and participating in active demethylation [76]. TET2 has been described as early and essential stage in somatic cell reprogramming preceding the induction of transcription at endogenous pluripotency loci. It is said to contribute to an epigenetic program that directs subsequent transcriptional induction at pluripotency loci during somatic cell reprogramming [77]. Adenosine deaminase (or adenosine aminohydrolase) ADA is located at 20q13.12 and is associated with severe immunodeficiency [78].

These genes and their expression in relevant tissues are shown in the Supplementary Material Figure 4. It will be interesting to track the evolution of these shared genes on induced pluripotent stem cell models, as cells develop into neuronal classes. Research along those lines is warranted [62]. Prostaglandin-endoperoxide synthase 2 PTGS2 or cyclooxygenase 2 COX2, is in 1q31.1. High-level induction of COX2 in mesenchymal-derived inflammatory cells suggests a role for COX2 in inflammatory conditions [79] and CNS-inflammatory pain hypersensitivity [80].

### 4.3. The Genes Common to Autism and All Subtypes

MECP2 and CREBBP were found to be shared pairwise with Autism and the above-mentioned disorders, but also present at the intersection set, taken across disorders. MECP2 expressed maximally in tissues related to emotion (amygdala) and memory (hippocampus) and tissues important for motor control (basal ganglia’s caudate and putamen regions, the substantia nigra, the cerebellum and cerebellar hemisphere, and the spinal cord) and for self-regulation (hypothalamus.) CREBBP was found to be maximally expressed at the cortex and frontal cortex, both of which are important for high-level cognitive and executive functions. Another important forkhead transcription factor FOXP1 was found to be maximally expressed in the basal ganglia’s nucleus accumbens, a structure important for developing striatal function and differentiation in medium spiny neurons from precursors to maturity [81–83]. CACNA1E was found maximally expressed across disorders in the anterior cingulate cortex, a region associated with impulse control, emotion and decision making, and previously known in connection to epilepsy, Autism, schizophrenia, and major depressive disorder [84–86].

Cluster analyses revealed that across all disorders under consideration, the cerebellum and cerebellar hemisphere had the maximal gene expression. Yet, different genes shared with the SFARI-Autism set contributed across disorders. MECP2 was maximally expressed in both the cerebellum and the cerebellar hemisphere in CP, Dystonia, OCD, Depression, PTSD and Lupus. In ADHD, MECP2 was maximally expressed in the cerebellum but the cerebellar hemisphere maximally expressed CREBBP. As mentioned, in PD, TSC2 was maximally expressed in both cerebellar tissues, while early onset had RAB39B and late onset had TET2 maximally expressed in both cerebellar tissues. Tourette’s had CHD2. Bipolar depression had SHANK2. Infantile Schizophrenia had ATP1A, and Schizophrenia had ATP2B2. These are shown in Supplementary Figures 4-6 along with other brain tissues and tissues important for cardiac, smooth, and skeletal muscles. The mixture of neuropsychiatric, neurological, and autoimmune disorders all had the cerebellar tissues with maximal gene expression of the genes shared with the SFARI-Autism set. These genes are thus bound to play an important role in motor control, coordination, initiation-termination, sequencing and adaptation, all critical components of basic building blocks to develop proper motor dynamics in social interactions. It is not surprising then that Autism has so many motor issues, as it sits squarely at the intersection of these neurological, neuropsychiatric, and autoimmune disorders. *Why are motor issues not seriously considered in Autism research and clinical practices*? Continuing to sideline the motor and motor sensing axes misses a superb opportunity to finally turn the science of Autism into a rigorous quantitative practice, beyond opinions or political agendas currently dominating the field and obfuscating important neurodevelopmental issues.

### 4.4. Implications of This Genomic Categorization for Treatment Selection in Autism

Approaching Autism as genotypically defined orderly subtypes may also be more humanely relevant to the affected individuals. Today, they receive recommendations for a “one-size-fits-all” behavioral-modification or conversion-therapy to reshape “socially inappropriate behaviors” without informing such treatments by brain-body physiological and medical issues. This approach disregards possible adverse effects linked to their genomic characteristics [87]. Indeed, despite advances in genetics, it has been reported that the current paradigm neglects the physiological phenotypes in favor of psychological constructs [33]. The literature reports that this model for Autism treatment selection promotes stigma, causes harm in the form of trauma, increases the person’s stress and ultimately results in PTSD [88]. Along those lines, there is a pool of genes linked to PTSD and depression overlapping with a subset of the SFARI-Autism data set, and very close to the Autism branch of the binary tree in Figure 5C. These genes may interact in ways that could increase the predisposition of the Autistic system to develop PTSD and depression, explaining the rise as well in suicidal ideation [51, 89–91].

The outcome of this work highlights the relevance of considering, when choosing treatments, the medical and physiological issues linked to the phenotypic characteristics that these genes forecast. This proposed approach contrasts with choosing treatments that exclusively focus on the social appropriateness criteria. The latter model has been said to lead to high societal cost [60, 92], to offer no future to the affected individuals and their families and has recently been shown to be polluted with conflict of interests and nonscientific practices [93, 94].

In the present study, I reasoned that stratifying Autism based on the genetic makeup of the diseases that today go on to receive the Autism clinical diagnosis, could help us in various ways. One was to find the diseases genetically closer to Autism itself (as defined by the SFARI genes.) Another was to leverage existing clinical information in other fields, amenable to create support and accommodations for the individuals affected by those disorders undergoing physiologically relevant treatments in other fields. Such accommodations could then be tailored to the autistic person, according to the phenotype that these genetic pools express for each of these other diseases of known etiology. Furthermore, since Autism today includes all these other disorders in its broad spectrum, utilizing the information that has already been verified (*e.g*., in the SFARI repository) would bring us a step closer to the Personalized Medicine approach, coined here *Precision Autism* (Figure 1B.)

It has been recently proposed that the behavioral definition of Autism, which recommends against stratifying the spectrum [13], feeds the Autism Industrial Complex (AIC) [92] and opens a behavioral diagnosis-to-treatment pipeline contributing to their claimed societal burden [60]. It is almost perverse to create a problem and sustain the problem by sidelining existing solutions, or alternative scientific routes, when the same model practiced over 40 years has not worked. It is as though to remain relevant and well-funded, that group steering autism research through the behavioral diagnostic-to-treatment pipeline, persists in neglecting the physiological issues.

The stratification of Autism revealed by the gene pool under consideration underscores the need to seriously consider the somatic-sensory-motor issues in the spectrum. This spectrum of disorders today includes diseases of known origins (*e.g*., Timothy Syndrome, SYNGAP1, SHANK3 deletion syndrome, Fragile X, Cerebral Palsy and Dystonia, among others) with life-threatening conditions that could seriously harm the affected child under the type of stress that a behavioral modification technique has been said to bring to their nervous systems [56, 87, 95].

This work revealed a compact set of top genes shared by SFARI-Autism and all diseases demonstrating that they too share tissues critical for *(i)* somatic-sensory-motor functioning, *(ii)* memory and cognition and *(iii)* systemic self-regulation. This compact set of genes for each of these functions in *(i-iii)* underly all critical physiological ingredients for social communication and smooth, well-coordinated actions. These basic functions are essential to all human autonomic, involuntary, and voluntary behaviors. As such, they should not be sidelined when recommending and selecting treatments for Autism. I provided a distance from each disorder to SFARI-Autism based on the genes’ expression on these 54 tissues defined by GETx, in the hopes of offering new ways to converge to truly personalized interventions that agree with the individual’s physiological phenotype and with the endophenotype of a genetically informed group.

I concluded from these analyses that the highly publicized exponential rate in prevalence reported by the US CDC surveillance network is a myth. This myth has been built by broadening and shifting the criteria over time and by allowing diseases of known etiology be part of the Autism spectrum. The increase in neurodevelopmental disabilities is real, as evidenced by the compact set of genes identified to be common to SFARI-Autism and all other diseases under consideration. All these genes play a fundamental role on the development of synapses via proteins that are necessary for channels functioning and neurotransmitters balance, neuronal differentiation, the formation of circuits and networks, *etc*., during neurodevelopment [96]. Yet, these disorders exist independent of Autism. Calling them Autism, under the current definition of inappropriate social behaviors may be doing more harm than benefit. The current model stigmatizes the affected individuals [60], their families and negatively impact the entire ecosystem inclusive of research, services and education, by promoting an erroneous perception of Autism as a behavioral issue [88]. By neglecting the physiology of the disorders that make up Autism today, the current approach skews the therapy recommendations for Autism in ways that may in fact harm their nascent nervous systems, induce trauma, and lower quality of life.

Given these results, I invite rethinking the epidemiology of autism spectrum disorders, to go beyond the behavioral diagnosis when surveying the spectrum to estimate prevalence. I also offer a new avenue to adapt the platform of Precision Medicine to Autism and disclose the implications of these results for the design of truly personalized therapies aimed at helping the affected individuals become an integral part of society.

### 4.5. The Importance of Reframing Autism under the Precision Medicine Paradigm

This notion of personalized medicine for Autism that I have proposed [25, 50], contrasts with the current behavioral diagnosis-to-treatment pipeline that discourages stratification of Autism and advocates for a general (one-size-fits-all) model of behavioral modification. Indeed, the last study from the U.S. National Academy of Science (NAS) considering how to educate autistic children, recommended that Autism shall not be stratified [13]. Since then, practice and services do not distinguish *e.g*., between a child with Cerebral Palsy and a child with ADHD. Both receive the Autism diagnosis, and both will receive a form of behavioral modification to reshape social behaviors in compliance with a set of social norms that bear no scientific empirical evidence for their recommendations. As mentioned, such imposed norms were never informed, in any way, by the nervous systems physiology [9]. The accreditation programs enabling such behavioral diagnoses and interventions in fact lack training on basic neuroscience^1^. In the US, these treatments will be administered at the school and the home, under a type of insurance coverage that other therapies do not have.

Our results show that contrary to the recommendations of the 2001 NAS study, such stratification is not only possible today, but more importantly, it is much needed to help guide and inform the design of new targeted therapies for Autism. Such new therapies could be truly personalized to address phenotypic features of the CNS, including tissues linked to self-regulating systemic structures, memory, cognition, and motor control. They would consider the physiology of numerous networks in the human brain-body complex, serving as the building blocks of all behaviors.

The methods used in this work are rather simple and parsimonious. They also rely on open access data sets. These sets are reliable and provide the grounds for reproducibility of this and related works [25, 50]. I encourage the community to stratify Autism into the appropriate phenotypes with capabilities, predispositions and needs causally linked to the genetic origins of each subtype. Continuing the blanket approach also misses three important revolutions of the 21^st^ Century: the genomic, the neuroscience and the wearables sensors revolution. The latter brings a level of precision to analyze continuous streams of behaviors beyond the limits of the naked eye, capable of automatically separating genetic-based disorders from natural, simple behaviors like walking and yet uncovering individual stochastic signatures of the person’s biorhythms with causal dynamics [44].

If we follow the medical and physiological scientific path, we will be able to advance Autism research, treatments, and services. But if we continue to follow the circular behaviorist approach, we will not make headways in identifying personalized targeted treatments. Worse yet, this antiquated approach, dating back to Skinner’s ideas of the 1950s, developed for research involving pigeons and rats, will continue to cause trauma to the individual in the spectrum. *Who came up with the notion of translating such methods to human children, without providing any validated scientific evidence that they would work in humas?* Such methods violate the natural autonomy of nascent nervous systems and go against the development of social agency [32, 97]. The current generation of adults that underwent such horror has informed us of this outcome. They have created the neurodiverse movement to alert researchers of the dangers of applying behaviorism to human babies in early intervention programs and throughout school age.

An alternative route to the current research paradigm in Autism is possible, by leveraging the work from other fields of science and engineering, and by stratifying the broad spectrum that otherwise purportedly keeps exponentially growing [1, 60]. Contrary to archaic recommendations from behaviorists [13], here I show that Autism can and should be stratified to take the first steps toward a paradigm shift toward *Precision Autism*.

## Supporting information

DisGeNet OCD genes shared with SFARI set

DisGeNet PD genes shared with SFARI set

DisGeNet PTSD genes shared with SFARI set

DisGeNet Schizophrenia genes shared with SFARI set

DisGeNet Tourette's genes shared with SFARI set

DisGeNet ADHD genes shared with SFARI set

DisGeNet Ataxia Dominant genes shared with SFARI set

DisGeNet Progressive cerebellar ataxia genes shared with SFARI set

DisGeNet Ataxia Recessive genes shared with SFARI set

DisGeNet X Ataxia genes shared with SFARI set

DisGeNet Bipolar Depression genes shared with SFARI set

DisGeNet CP genes shared with SFARI set

DisGeNet Depression genes shared with SFARI set

DisGeNet Dystonia genes shared with SFARI set

DisGeNet Fragile X genes shared with SARI set

DisGeNet FXTAS genes shared with SFARI set

## Supplementary Materials

The following are available online at www.mdpi.com/xxx/s1, Figure S1: Colormaps of top 20 genes expressed in 54 tissues at the intersection of SFARI-Autism and other disorders-I, Figure S2: Colormaps of top 20 genes expressed in 54 tissues at the intersection of SFARI-Autism and other disorders-II, Figure S3: Colormaps of top 20 genes expressed in 54 tissues at the intersection of SFARI-Autism and another disorders-III, Figure S4-S6 colormaps of genes shared between SFARI-Autism and other disorders, maximally expressed in brain tissues and tissues linked to cardiac, smooth and skeletal muscles across disorders. Text files list the genes at the intersection of SFARI-Autism and other disorders.

## Author Contributions

Conceptualization, methodology, software, validation, formal analysis, investigation, resources, data curation, writing, visualization, supervision, project administration, funding acquisition, E.B.T.

## Institutional Review Board Statement

Not applicable. Although the study involves de-identified data from human participants, these data is open access and required no IRB protocol/approval.

## Data Availability Statement

Data supporting the results can be found at https://www.sfari.org/resource/sfari-gene/, https://gtexportal.org/home/ and https://www.disgenet.org/.

## Acknowledgments

I thank the families that participated in the genetic studies providing the data for SFARI, GTEx and DisGeNet and the supporters of OMIM, as well as the many researchers that upkeep these data repositories.

## Footnotes

1 https://accreditation.abainternational.org/apply/accreditation-standards.aspx https://www.wpspublish.com/ados-2-autism-diagnostic-observation-schedule-second-edition

