## Supplementary figures and images for "Precision Autism: Genomic Stratification of Disorders Making Up the Broad Spectrum May Demystify its “Epidemic Rates”"

### Supplementary Figure 1

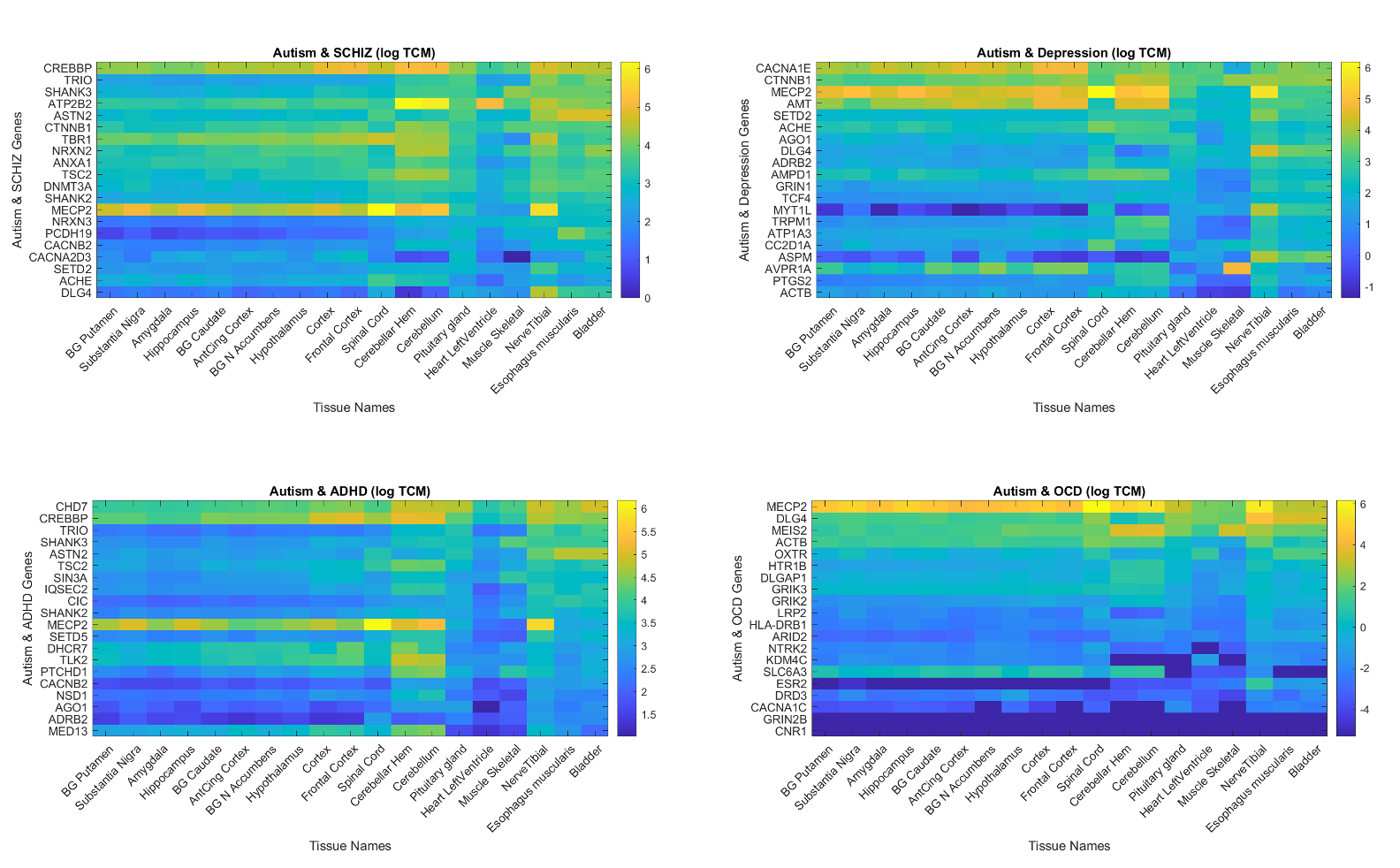

### Supplementary Figure 2

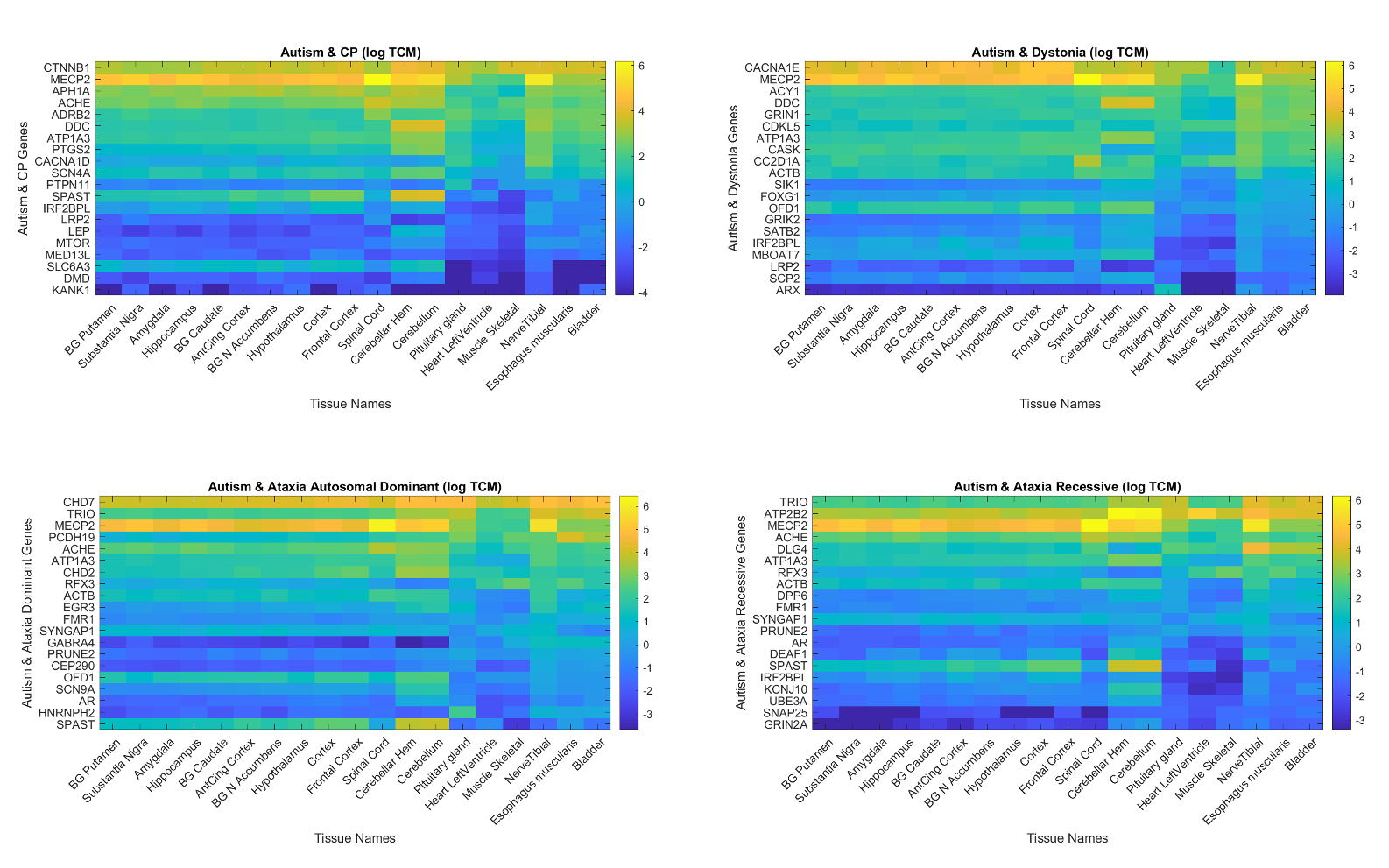

### Supplementary Figure 3

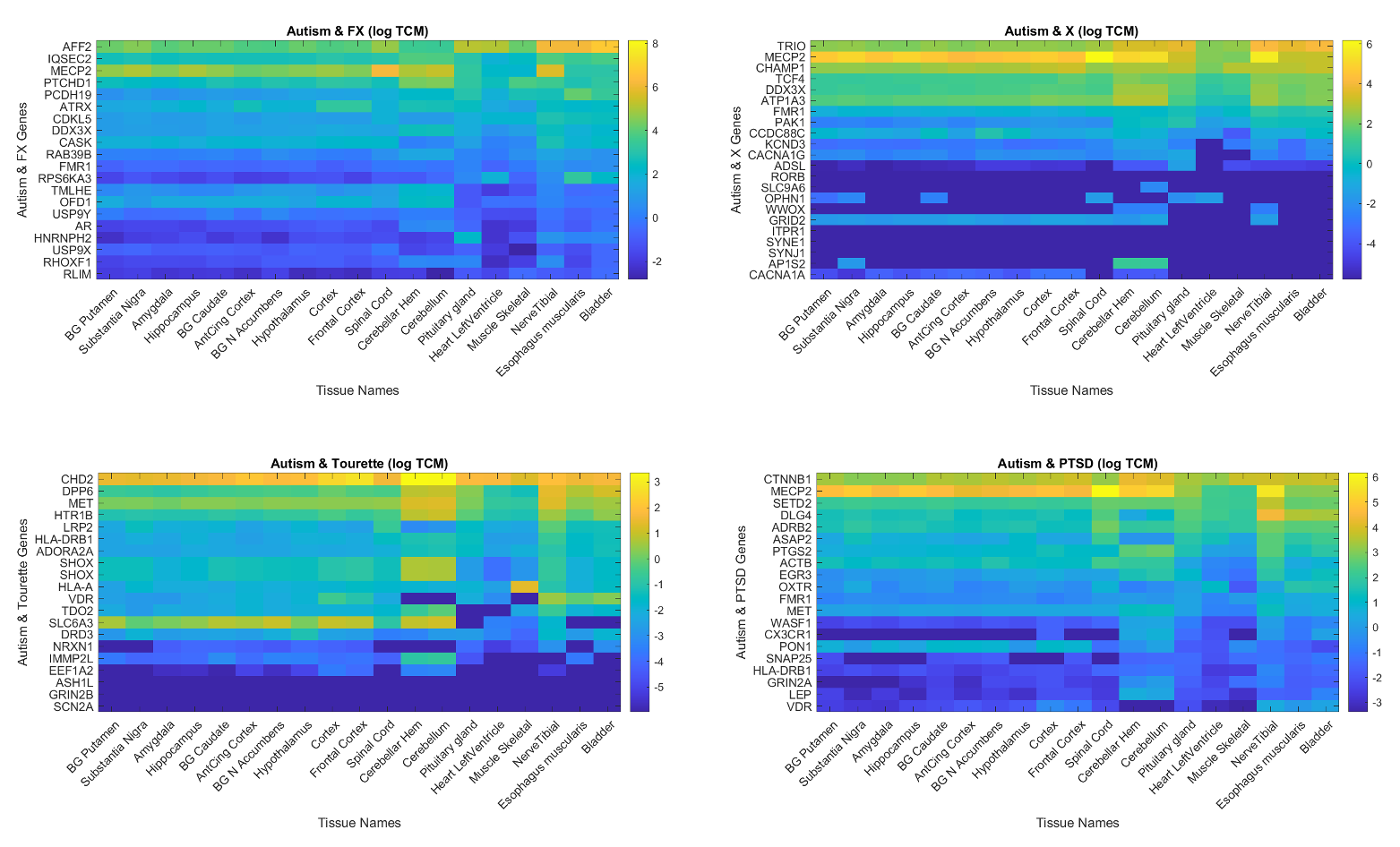

### Supplementary Figure 4

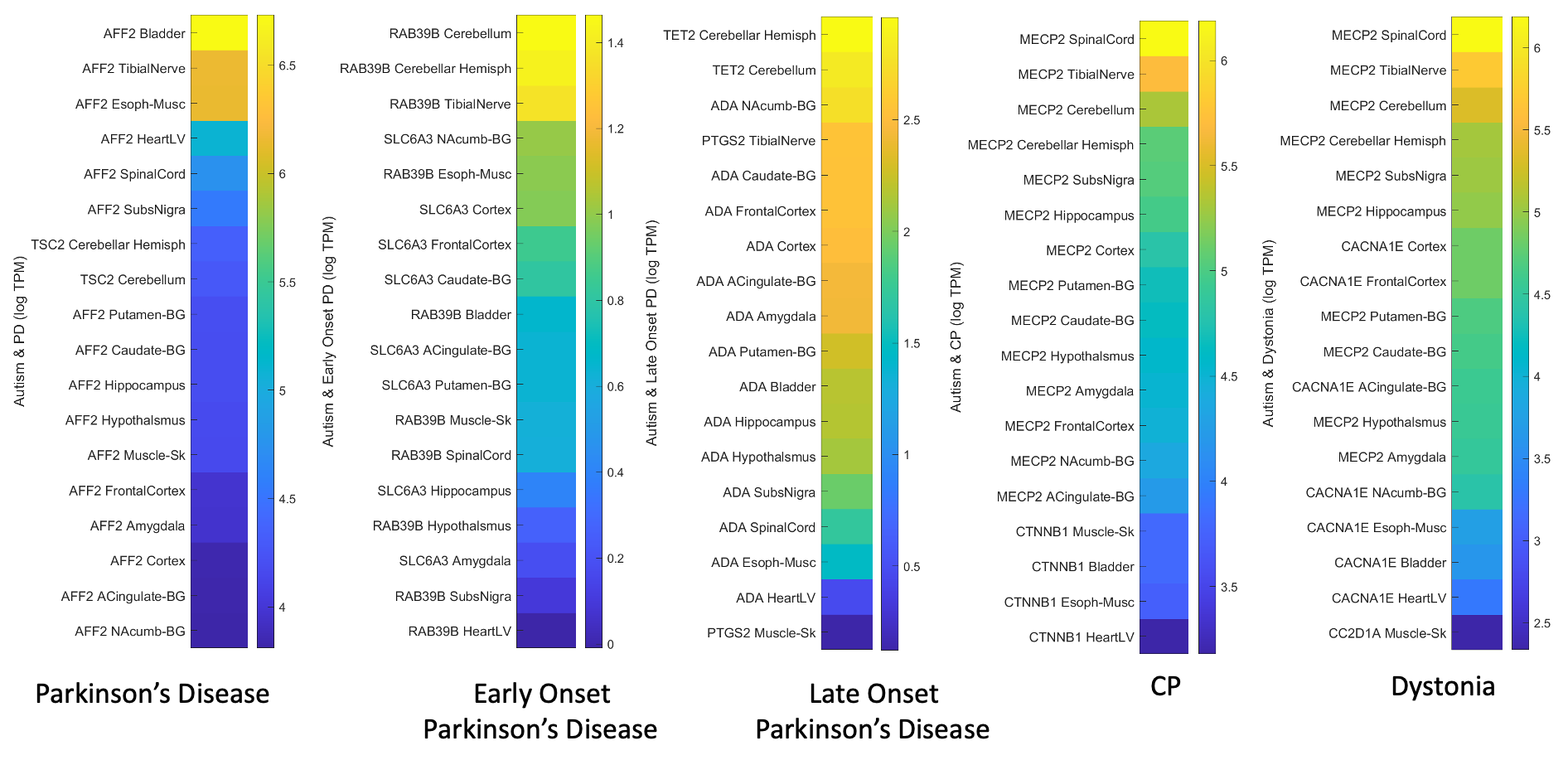

### Supplementary Figure 5

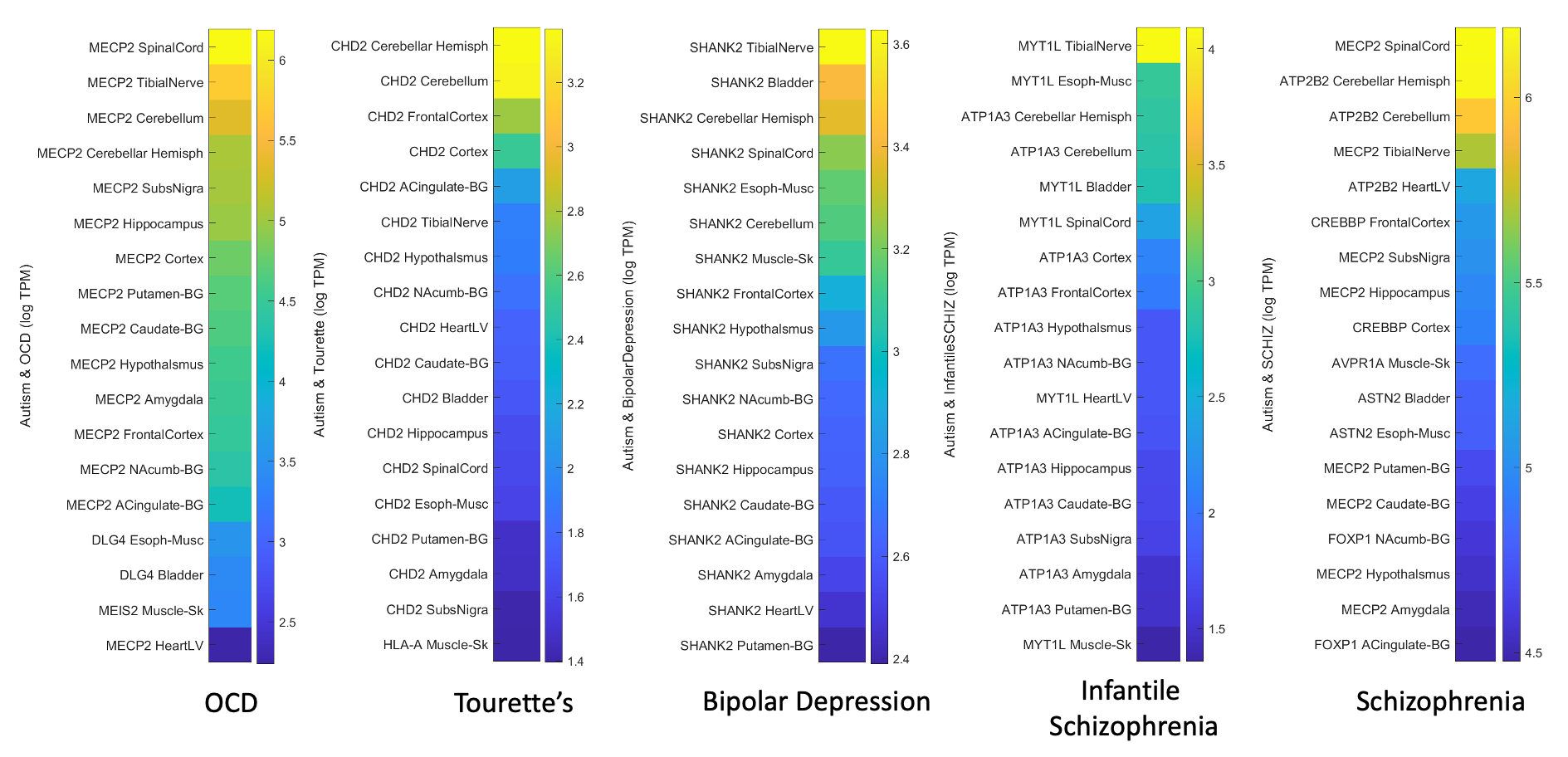

### Supplementary Figure 6

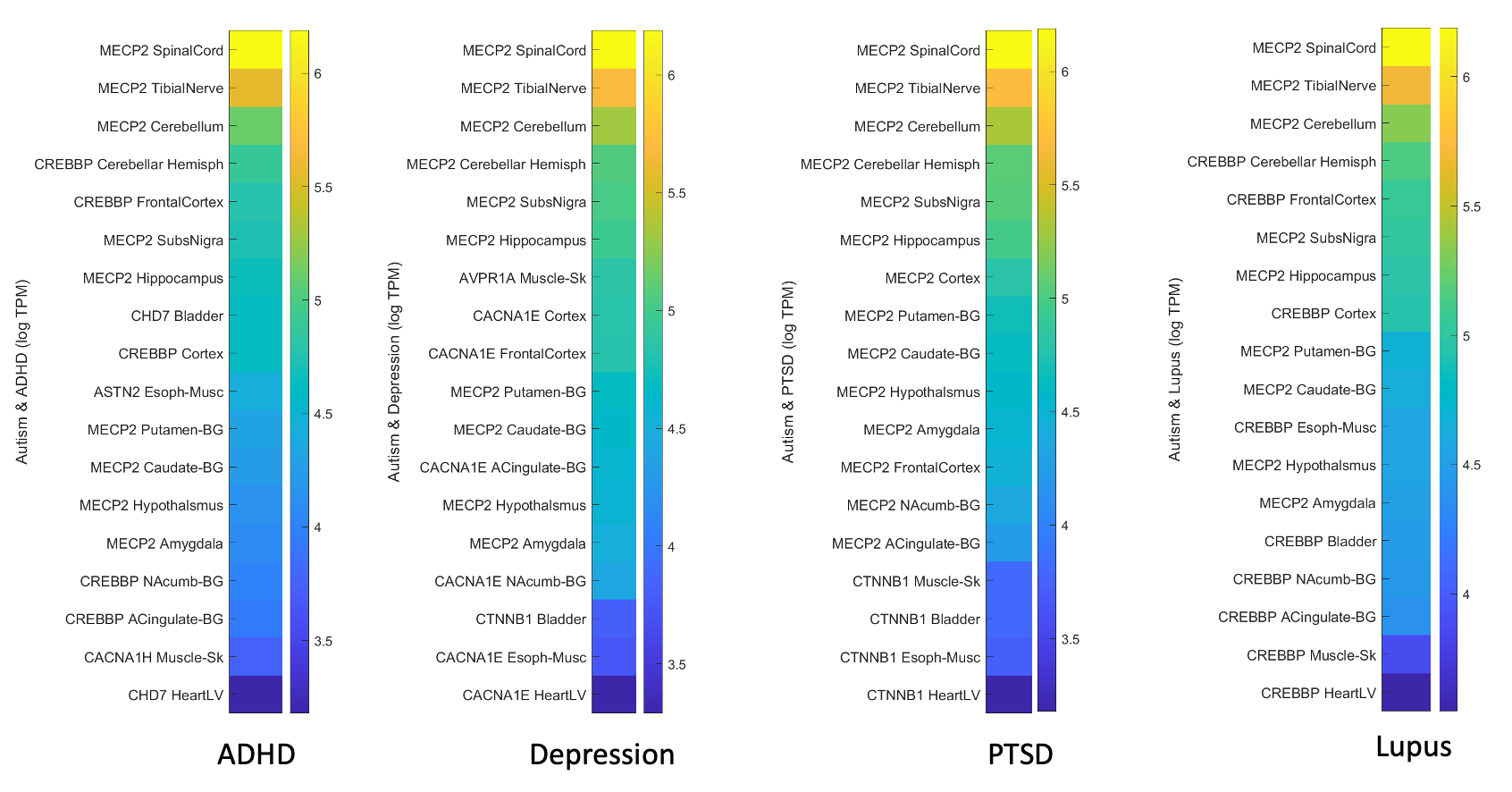
